# Efficient differentiation of human primordial germ cells through geometric control reveals a key role for NODAL signaling

**DOI:** 10.1101/2021.08.04.455129

**Authors:** Kyoung Jo, Seth Teague, Bohan Chen, Hina Aftab Khan, Emily Freeburne, Hunter Li, Bolin Li, Jason Spence, Idse Heemskerk

**Affiliations:** Department of Cell and Developmental Biology, University of Michigan Medical School, Ann Arbor, Michigan; Department of Biomedical Engineering, University of Michigan, Ann Arbor, Michigan; Center for Organogenesis, University of Michigan Medical School, Ann Arbor, Michigan; Department of Internal Medicine, Gastroenterology, University of Michigan Medical School, Ann Arbor, Michigan; Department of Physics, University of Michigan, Ann Arbor, Michigan

## Abstract

Human primordial germ cells (hPGCs) form around the time of implantation and are the precursors of eggs and sperm. Many aspects of hPGC specification remain poorly understood. Here we show that micropatterned human pluripotent stem cells (hPSCs) treated with BMP4 give rise to hPGC-like cells (hPGCLC) and use these as a quantitatively reproducible and simple *in vitro* model to interrogate this important developmental event. We characterize micropatterned hPSCs up to 96h for the first time and show that hPGCLC populations are stable and continue to mature. By perturbing signaling during hPGCLC differentiation, we identify a previously unappreciated role for NODAL signaling and find that the relative timing and duration of BMP and NODAL signaling are critical parameters controlling the number of hPGCLCs. We formulate a mathematical model for a network of cross-repressive fates driven by NODAL and BMP signaling which predicts the measured fate patterns after signaling perturbations. Finally, we show that hPSC colony size dictates the efficiency of hPGCLC specification, which led us to dramatically improve the efficiency of hPGCLC differentiation over current protocols.

## INTRODUCTION

Primordial germ cells (PGC) are the first step in specification of the germline, the unique lineage through which genetic material is passed on to the next generation and potentially the key to understanding totipotency. Germline defects underlie numerous human diseases, most notably infertility(Chen, Gell, et al. 2017). Understanding PGC specification is therefore critical both for our fundamental understanding of human development, and for its practical implications in disease. Yet, human germline specification remains an elusive process. Until recently, mammalian PGC specification was predominantly studied in mice. However, significant interspecies differences in PGC specification have been documented, in particular between rodents and primates but possibly also within the primates(Kojima et al. 2017; Kobayashi et al. 2017; Kobayashi and Surani 2018). Because there is limited access to pre-implantation human embryos(Hancock et al. 2021) and it is not admissible to study post-implantation human embryos, non-human primates and human pluripotent stem cell (hPSC)-based models of PGC differentiation have played a key role in advancing our understanding of this process. *In vitro* differentiation of PGC-like cells (PGCLCs) from human pluripotent stem cells (hPSCs) has been essential in revealing key aspects of hPGCLC specification such as the transcription factor network involving SOX17, PRDM1, and TFAP2C (Irie et al. 2015; Chen et al. 2019; Kojima et al. 2017). However, like many directed differentiation processes, hPSC-derived PGC specification is variable from batch to batch and line to line(Chen et al. 2019). This makes it difficult to systematically and quantitatively study how and where PGCLCs arise in cell culture models. Although major progress has been made, much about germ cell specification remains poorly understood. For example, it is unknown whether human PGCs derive from the primitive streak like in mouse and pig, or from the amnion like in cynomolgous monkeys(Kobayashi et al. 2017; Lawson et al. 1999; Sasaki et al. 2016). It also remains unclear what the precise cell signaling requirements are that separate PGC specification from amnion on the one hand and mesendoderm on the other hand.

Here we used micropatterned hPSCs as a quantitatively reproducible system that allowed systematic interrogation of hPGCLC specification at single cell resolution. Micropatterning enables spatial restriction of substrate adhesion by cells to control colony size and shape. Micropatterned human embryonic stem cells treated with BMP4 for 42-48h are a model system of human gastrulation, generating all 3 germ layers in concentric rings surrounded by another ring of extraembryonic-like cells(Warmflash et al. 2014). The inner domain consists of ectodermal or pluripotent cells depending on the differentiation media(Chhabra et al. 2019). Surrounding the inner domain is a ring of cells expressing primitive streak markers such as BRA and EOMES. The outer ring of cells on the colony edge was initially thought to be trophectoderm-like due to its expression of CDX2 in the absence of BRA but was later found to have features of both amnion and trophectoderm(Chhabra and Warmflash 2021; Minn et al. 2020).

A final ring of SOX17 positive cells, roughly positioned between the extraembryonic cells and primitive-streak-like cells was originally thought to be endoderm. However, these SOX17+ cells do not express the definitive endoderm marker FOXA2+(Martyn, Siggia, and Brivanlou 2019). Moreover, they are positioned close to the colony edge where BMP signaling is high. In contrast, directed endoderm differentiation from hPSCs is improved by BMP inhibition(Loh et al. 2014) and in the mouse embryo endoderm is thought to arise from the anterior streak where BMP is low(Nowotschin, Hadjantonakis, and Campbell 2019). Here we further investigate the identity of each of the cell types and report that this puzzle is resolved by the finding that the SOX17+ cells juxtaposed with the extraembryonic tissue at 42h are not endoderm but PGCLCs, confirming what was also recently reported in(Minn et al. 2020). Although SOX17+ uniquely marks endoderm in the mouse, it is well known to be expressed in primate PGCLCs and the location of the PGCLCs in our system is consistent with mouse development, where PGCs arise in posterior streak on the interface with the extraembryonic tissue in a BMP-dependent manner.

We developed improved quantitative analysis of immunofluorescence (IF) data at the single cell level, based on a 3D image analysis pipeline integrating deep-learning based segmentation. This enabled accurate assessment of the molecular signatures, spatial distributions, and sizes of cell populations. We combined this with scRNA-seq to confirm PGCLC identity and further found evidence of amnionic ectoderm identity of the outer ring. By carrying out a temporal analysis up to 96h we find that PGCLC populations persist and mature during this time window.

After confirming PGCLC specification, we carried out pharmacological and genetic perturbations to provide more insight into the underlying signaling involved in this process. Although a requirement for NODAL in mouse PGC differentiation was demonstrated(Senft, Costello, et al. 2018; Senft et al. 2019; Mulas, Kalkan, and Smith 2017), directed differentiation of human PGCLCs has focused on the BMP and WNT and the precise roles and interplay of these pathways remains unclear(Hancock et al. 2021; Kobayashi et al. 2017). We confirm a requirement for continuous BMP signaling for the first two days of hPGCLC differentiation but find that WNT signaling is only required within a short time window from 12-24h and provide evidence that the primary role for WNT is to induce NODAL. We show that NODAL is required for hPGCLC induction and that exogenous stimulation of the NODAL pathway can rescue PGCLC induction when WNT is inhibited. We find that the timing and duration of NODAL are critical in deciding between amnion-like, PGCLC and primitive streak-like fates. In addition, we find FGF/ERK signaling is essential throughout differentiation.

Finally, we investigate how PGCLC differentiation depends on colony size and find that by optimizing colony size, we can generate PGCLCs with efficiencies of ~50% using BMP4 treatment alone. When 12h of pre-differentiation to an incipient mesoderm like state (iMeLC) is included, as is typical in directed PGCLC differentiation, this number goes up to 70%, compared to other literature reporting 20-30% efficiency(Sebastiano et al. 2021; Sasaki et al. 2015).

## RESULTS

### PGCLCs form on the interface between extra-embryonic and primitive streak-like cells

Upon treatment with BMP4, at least 4 distinct cell fates arise in concentric rings by 42h in micropatterned hPSC colonies with 500-1000um diameter(Warmflash et al. 2014). Cells expressing SOX17 have repeatedly been identified as endoderm (Warmflash et al. 2014; Martyn, Siggia, and Brivanlou 2019). Curiously however, we found that these cells do not express the definitive endoderm marker FOXA2 (Fig 1A). In primates, SOX17 does not only mark definitive endoderm but also primordial germ cells (PGCs), which moreover form close to the interface of the posterior epiblast and amnion, with their precise origin in human still a point of debate(Saitou 2021; Hancock et al. 2021). This suggests these cells could be PGC-like cells (PGCLCs) instead. To test this idea, we used immunofluorescence to visualize the marker genes TFAP2C, PRDM1 and NANOG which in combination are known to uniquely mark PGCLCs(Tyser et al. 2021; Yang et al. 2021). This confirmed our hypothesis and showed the reproducible presence of PGCLCs (Fig. 1B), positioned between ISL1+ extraembryonic cells and EOMES+/TBXT+ PS-like cells, (SI Fig 1 A-D).

**Figure 1:**
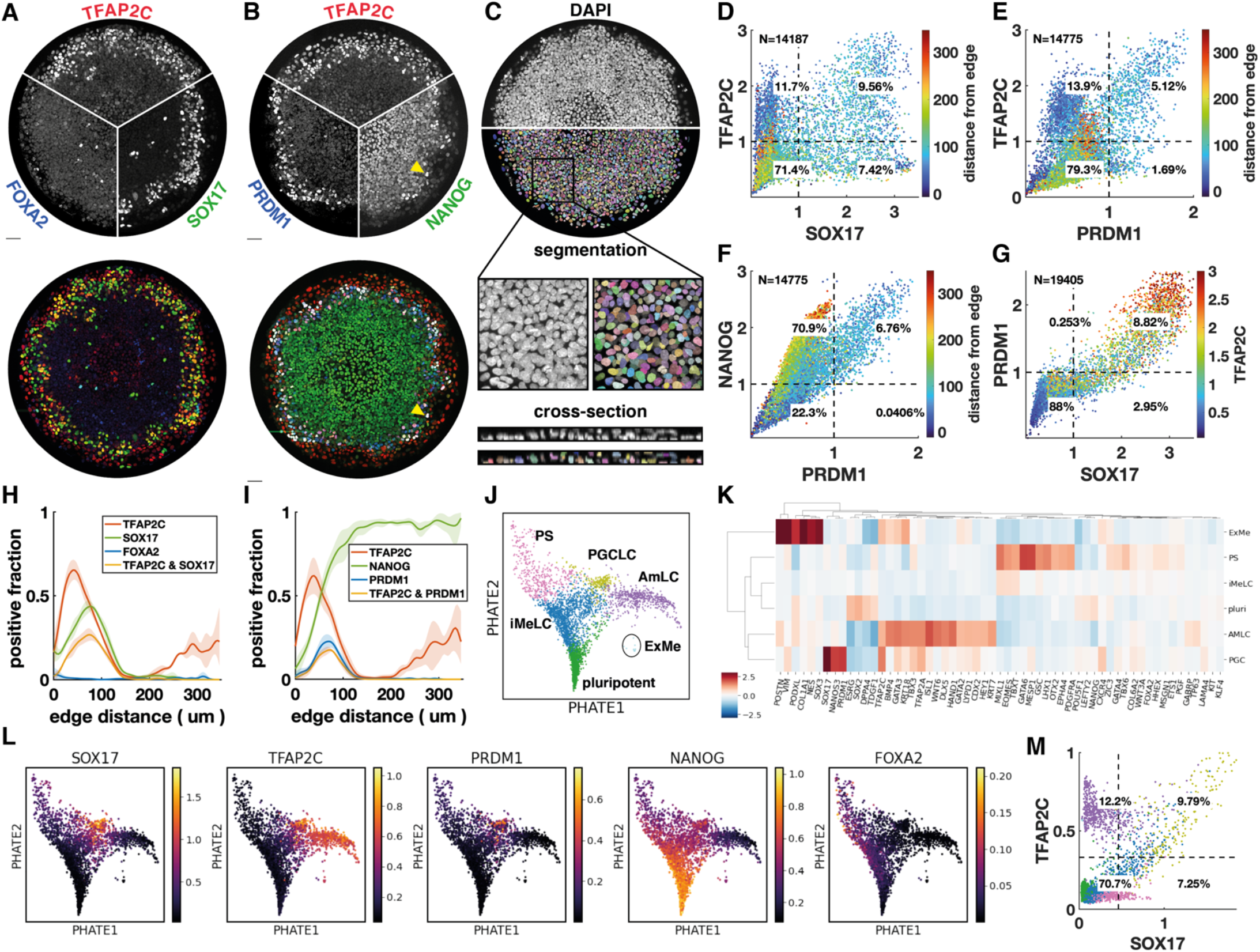
PGCLCs form on the interface between extra-embryonic and primitive streak-like cells. **a-b)** Immunofluorescence for different marker genes (maximal intensity projection along z). Yellow arrowhead in b points to higher NANOG expression in PGCLCs than pluripotent cells in the colony center. **c)** Segmentation of nuclei based on DAPI staining. **d-f)** Scatter plots of marker expression colored for radial position, normalized to threshold and log(1+x) transformed, d corresponds to a; e,f correspond to b. **g)** scatter plot of PRDM1 vs. SOX17 colored for TFAP2C. **h-i)** spatial distribution of positive cells, dark lines represent the mean kernel density estimate of the positive fraction over 4 colonies, colored bands represent the standard deviation. **j)** Clusters generated by Louvain. **k)** Heatmap of differential expression between clusters (average z-scores) of genes associated with gastrulation. **l)** PHATE visualization of scRNA-seq data showing expression of markers used in b-i. **m)** Scatterplot of TFAP2C vs. SOX17 from denoised scRNA-seq data, with colors matching clusters in k. Scalebars 50um. All colonies are 700um diameter.

To quantify the relationship between these markers from the immunofluorescence data, we developed a 3D image analysis pipeline based on machine learning to handle multiple overlapping cell layers and automatically determine if cells express or co-express specific markers (see methods). Segmentation (Fig 1C) allowed us to identify both expression and relative level of expression (within an individual image) in order to generate single cell scatter plots of protein expression that show different co-expressing groups of cells, in a manner similar to data from flow cytometry (Figure 1 D-G).

Across multiple experiments and cell lines (SI Fig 3), we found ~10-20% of cells to be SOX17+ with 50-60% of those also expressing TFAP2C (Fig. 1D). About 5-10% of cells were found to be PRDM1+ which were mostly TFAP2C+ (Fig. 1E), and all NANOG+ (Fig. 1F). Moreover, PRDM1+ were nearly all SOX17+ (Fig. 1G, SI Fig. 1EF), consistent with previous literature showing that SOX17 is upstream of PRDM1(Kojima et al. 2017). Most SOX17+ PRDM1+ expressed higher TFAP2C than SOX17+PRDM1-(Fig 1G), and of SOX17+TFAP2C+, 80% were PRDM1+ (SI Fig. 1G,E-H). Thus PRDM1+TFAP2C+ implies PRDM1+TFAP2C+SOX17+NANOG+ and provides a conservative estimate of the PGCLC population while SOX17+TFAP2C+ provides a similar but slightly higher estimate. Here, we will use both combinations to quantify the PGCLC population. We observed that both TFAP2C and NANOG levels in PGCLCs are higher than in non-PGCLCs (Fig 1B,G), suggesting a positive feedback in the co-expression of these factors.

After identifying populations by thresholding markers, we visualized spatial patterning as the fraction of cells positive for a marker at some radius (Fig 1H-I). We found this to be a substantial improvement over average intensity profiles that have so far been used in studying micropatterned hPSCs (SI Fig. 1HI), because the relative magnitude of the markers in the graph becomes meaningful and it eliminates the effect of background when positive cell populations are small (see FOXA2 in SI Fig 1H). Moreover, it allows simple visualization of the spatial of marker combinations like TFAP2C & SOX17 (Fig. 1HI).

We repeated quantitative analysis of immunofluorescence for PGC markers with 4 different hPSC lines, both male and female (SI Fig. 3). We found that all these form similar patterns although with some variability in the fraction of PGCLCs.

### scRNA-seq confirms PGCLC identity and shows extra-embryonic cells resemble amnion

To further understand the identity of both the SOX17+ cells and other cells within the micropatterned hPSCs we performed scRNA-seq and visualized our data using PHATE (Moon et al. 2019). This reproduced the known gene expression domains and organized them in a lineage tree-like layout with SOX2+ pluripotent cells at the bottom, a TBXT+ primitive streak-like branch on the left, a ISL1+ branch on the right, with a group of SOX17+ cells in between these two branches (SI Fig. 2AB). Diffusion components showed the SOX17+ cells more clearly as a third branch (SI Fig. 2C).

To systematically evaluate gene expression in PGCLCs, we performed clustering using Louvain, which yielded six clusters (Fig. 1J). We performed differential expression analysis to identify marker genes for each cluster, both for all genes and within a subset of marker genes relevant for gastrulation (SI tables 1–3). In addition, we found it instructive to visualize differential expression in a subset of marker genes that are commonly used to identify cell fate during gastrulation (Fig. 1K). As expected, four of the clusters found by Louvain corresponded roughly to the cell groups identified using immunofluorescence: pluripotent, PGCLC, extraembryonic, and PS-like. Confirming the immunofluorescence, PGCLCs are marked by highly enriched expression of SOX17, PRDM1, TFAP2C, while FOXA2 expression was low and not in the same cells that expressed high SOX17 (Fig. 1KL). In contrast to IF data, NANOG levels in PGCLC are significant but lower than in pluripotent cells, suggesting posttranscriptional regulation to explain why NANOG protein levels are higher in PGCLCs. As expected, the PGCLC cluster also showed high expression of NANOS3, which is known to be uniquely expressed in PGCLCs (Fig. 1K, SI Fig. 2D).

The identity of the outer ring of cells has been a source of debate and is important in the context of PGCLC induction because PGCs have been found to derive from the amnion in cynomolgous monkeys, while their origin in human remains unclear(Saitou 2021; Hancock et al. 2021). We argue that these cells are amnion-like and named them AmLC.

The outer cells were previously found to express markers of both trophectoderm (TE)) including CDX2, GATA3, TP63, TBX3, and KRT7, but also genes associated with amnion such as TFAP2A, with little consensus on which genes specifically mark human AM in vivo (Chhabra and Warmflash 2021; Minn et al. 2020) (Chen et al. 2019; Sasaki et al. 2016; Knöfler et al. 2019). Recently, ISL1 and BMP4 were established as key amniotic genes and GABRP and WNT6 as additional amnion markers(Yang et al. 2021). We found the cells of the outer ring express all the above (Fig. 1K, SI Fig. 2D), raising the question of whether this is a state between amnion and trophectoderm that is an artefact of the in vitro system. However, there is no published complete expression profile of both amnion and trophectoderm from a single human or even non-human primate embryo to validate the presumed markers and compare the two tissues. Nevertheless, we found that there may be in vivo transcriptome data for human amnion from the CS7 human gastrula in (Tyser et al. 2021). They found is a cluster of cells labeled non-neural ectoderm, which was noted by the authors to be also consistent with amnion. Importantly, these cells express all the markers mentioned above (SI Fig. 2E). We quantify the similarity between our clusters and those in the human gastrula dataset by cross-correlating gene expression (SI Fig 2F). Strikingly, the strongest correlation between any two clusters was between our AmLC and the cells labeled non-neural ectoderm in Tyser et al. The placement in UMAP of this cluster, branching off between epiblast and primitive streak close to the PGCs is also very similar to what we see in our PHATE plots (Tyser Fig. 2A). Furthermore, due to the way the sample was dissected it was unlikely to contain trophectoderm. All of this suggests that the outer ring on micropatterns are amnionic ectoderm, and that the expression of several markers that were thought to be trophectodermal is a feature of human amniotic ectoderm. However, until unambiguous in vivo data is published comparing amnion, trophectoderm and non-neural ectoderm, or functional data can be obtained, we cannot be completely certain of the identity of this cell population.

**Figure 2:**
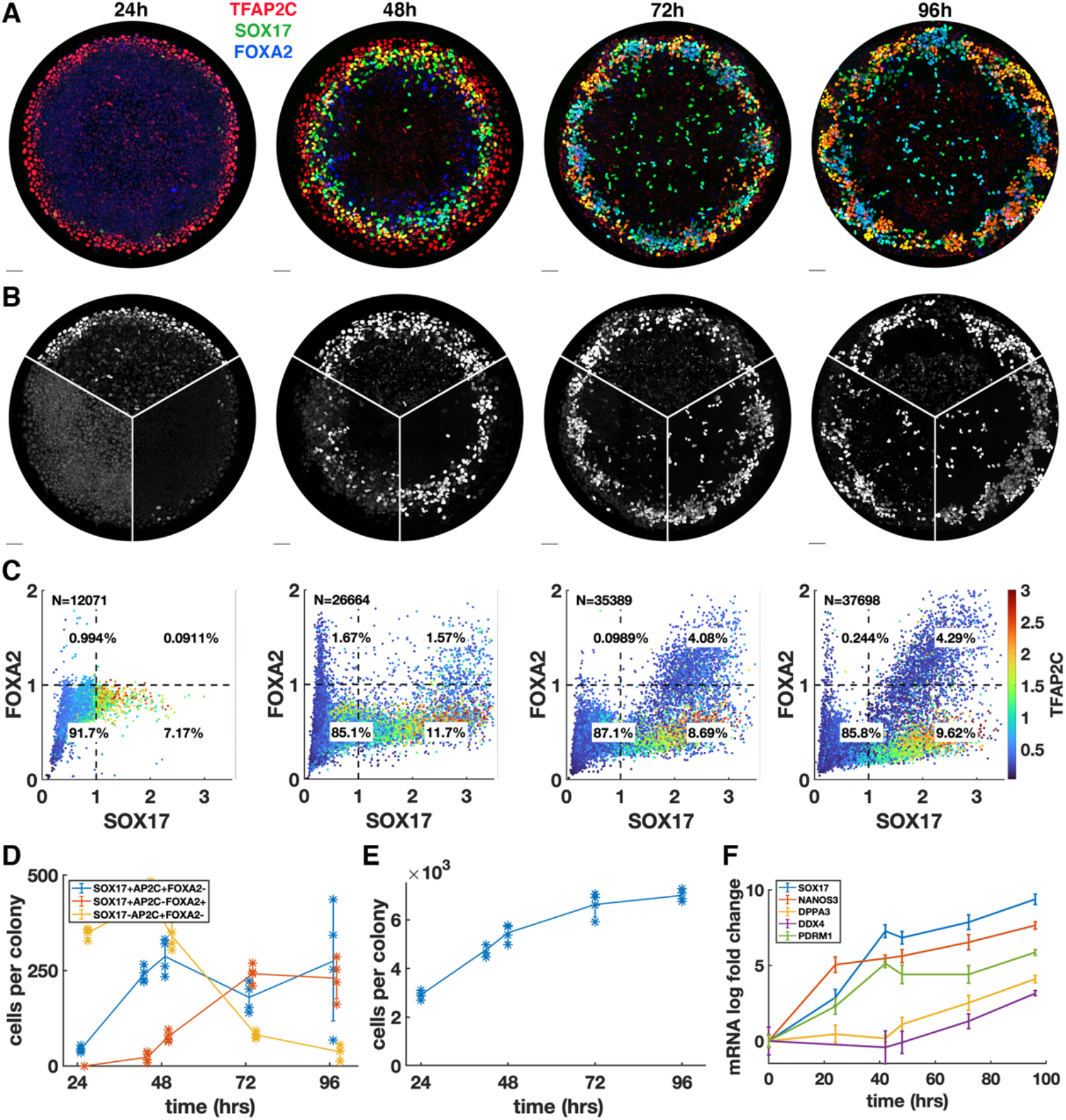
PGCs are specified by 42h but continue to mature while endoderm arises between 42h and 72h. **a-b)** Immunofluorescence over time showing stable PGC population and later emergence of endoderm **c)** Quantification of marker expression at different times showing the emergence of endoderm starting at 48h. **d)** Absolute numbers of cell expressing marker combinations corresponding to endoderm (red, SOX17+AP2C-FOXA2+) PGCLCs (blue, SOX17+AP2C+FOXA2-) and SOX17-AP2C+FOXA2-(yellow). **e)** average cell number per colony over time. **f)** qPCR data for PGC markers over time. DAPI stainings corresponding to a-b) are shown in SI Fig. 4. Scalebar 50um.

A fifth cluster was identified between the pluripotent cells and differentiated fate, expressing intermediate levels of both primitive streak and pluripotency markers. This appears to be a transitionary state. In this context we decided to name this intermediate state incipient mesoderm-like (iMeLC) as is used in directed differention of PGCLCs(Sasaki et al. 2015).

The sixth and final cluster of cells was very clearly distinct from other cells in the scRNA-seq data; however very few cells were captured in this cluster suggesting these cells are rare within micropatterns. This cluster was enriched for ANXA1, POSTN, VIM and PODXL (Fig. 1K, SI tables) suggesting a yolk sac-mesoderm or extraembryonic mesenchyme identity. However, expected expression of GATA6 is missing. Moreover, it would be very surprising to find extra-embryonic mesenchyme, which is thought to derive from the hypoblast, a tissue that is absent from our in vitro model. Correlation with CS7 gastrula data nevertheless did show strong correlation between this cluster and the YS mesoderm (SI Fig.2E) so we decided to tentatively label it extraembryonic mesoderm (ExMe). Given the small number of cells expression data for this cluster is very noisy and future investigation will have to confirm the consistent presence, identity, and origin of these cells.

### Immunofluorescence and scRNA-seq reveal similar quantitative gene relationships

We asked whether the gene expression relationships found using immunofluorescence in Fig. 1G-J could also be recovered when considering the same 2 genes in the scRNA-seq data, and whether the clusters obtained based on 2 markers correspond to the clusters identified in the full scRNA-seq dataset. While raw scRNA-seq data is too noisy to directly relate expression of two genes (SI Fig 1J), after denoising using MAGIC(van Dijk et al. 2018), clear patterns emerged (Fig. 1M). Thresholding using the same procedure used for IF produced nearly identical proportions of cells expressing SOX17 and/or TFAP2C with most of the cells in the SOX17+TFAP2C+ quadrant belonging to the PGCLC cluster found by Louvain, demonstrating consistency between the data types and clustering procedures. Fig. 1M further suggests that a significant fraction of SOX17+TFAP2C- cells belong to the iMeLC cluster and are on their way to becoming PGCLCs.

It remains unclear whether human PGCs derive from amnion or posterior epiblast. Moreover, PGCs are thought to go through an incipient mesodermal state transiently expressing low levels of primitive streak (PS) markers. We therefore also looked at co-expression of PGC markers with the PS markers EOMES and TBXT, and the amnion marker ISL1 (SI Fig 1).

EOMES is required to induce SOX17 during human PGCLC specification but is then rapidly downregulated(Kojima et al. 2017; Chen, Liu, et al. 2017). In the mouse EOMES is not directly required in PGCLC induction, but it is required for EMT in gastrulation and specification of endoderm and cardiac mesoderm(Senft, Bikoff, et al. 2018; Costello et al. 2011; Tosic et al. 2019; Arnold et al. 2008). EOMES knockout hPSCs suggest that these functions in gastrulation are conserved in human (Teo et al. 2011; Pfeiffer et al. 2018). While in the overlay image it appears as if ISL1 forms a shape boundary and has little overlap with EOMES and SOX17 (SI Fig. 1A), the individual images and quantification (SI Fig 1A,K) show low EOMES and ISL1 expression in SOX17+ cells. The scatter plot shows a striking inverse relationship between ISL1 and EOMES, suggesting mutual repression, with EOMES^low^ ISL1^low^ SOX17+ cells connecting EOMES+ISL1-cells to EOMES-ISL+. This is consistent with a requirement for, but subsequent suppression of, EOMES in PGCLCs and places the gene expression profile of PGCLCs intermediate between amnion and primitive streak. Similarly, we found that TFAP2C+ cells that co-express TBXT are mostly PRDM1+ and show a strong correlation between TFAP2C and PRDM1 within this cluster (SI Fig. 1C,M).

Strikingly, the relationships produced by plotting scRNA-seq data for the same genes look very similar, in particular for ISL1 vs. EOMES (SI Fig. 1L,N). Some differences do exist, e.g. the gap between TFAP2C+ and TFAP2C- along the TFAP2C axis in the scRNA-seq data, which is not present in the IF data (Fig. 1M, SI Fig. 1N). There are several possible explanations for this: it may be a difference between RNA and protein levels, the gap could be an artifact of the denoising algorithm on the single cell RNA-seq data, or the lack of a gap could be due to noise in the IF data. Nevertheless, these data show that consistent quantitative relationships can be recovered from these different types of data which may inform mathematical models for the underlying gene regulatory networks.

### PGCs are specified by 42h but continue to mature through 96h, while endoderm arises between 42h and 72h

We next asked whether the PGCLC population would persist and continue to develop past the 48h during which BMP4-treated micropatterned colonies have so far been studied. In addition, we wanted to know whether definitive endoderm arises later. Therefore, we quantified the time course of TFAP2C, SOX17 and FOXA2 with 24-hour resolution up to 96 hours (Fig 2) and also included 42 hours in the analysis because this is the end point used in other experiments (Fig. 2D-F). Immunofluorescence immediately revealed significant changes between 48h and 72h with a striking pattern of alternating clusters of PGCLCs (SOX17+TFAP2C+) and endoderm (SOX17+FOXA2+) appearing around the perimeter by 72h.

Our quantification showed that in contrast to 42h (Fig 1A) small numbers of FOXA2+ cells, both SOX17+ and SOX17-, emerge at 48h followed by a large increase in FOXA2+SOX17+ cells between 48h and 72h, while FOXA2+SOX17-were no longer present at 72h (Fig 2CD). Quantitative analysis confirmed FOXA2+SOX17+ are TFAP2C- while FOXA2-SOX17+ are TFAP2C+ (Fig 2C), consistent with PGC and endodermal populations. We conclude that endoderm is specified between 42h and 72h. In contrast PGCs may be fully specified before 42h and do not appear to proliferate after that since PGC numbers are stable between 42h and 72h. Between 72h and 96h we observed no significant changes in either cell population.

Since the 72h and 96h time points have not been looked at before, we also looked at overall growth and morphology of the colonies over time. We found that the growth rate is gradually decreasing from 60% increase in cell number from 24-48h, a 38% from 48-72h and 5% from 72-96h (Fig 2E and SI Fig 4A). Because of the continued growth but stable PGC population, the percentage of PGCs goes down over time which is reflected in the spatial distributions (SI Fig. 4B). Looking at the 3D structure, we found that colonies become significantly thicker between 48h and 72h forming either a multilayered structure or pseudostratified epithelium (SI Fig. 4CD). Peripheral endodermal and PGC clusters appear near the top of the colony while scattered SOX17+ cells throughout the center of the colony are on the bottom of the colony. From 72h to 96h the colony undergoes a slight morphological change, expanding outward beyond the borders of the micropattern and thinning in the colony center while the positioning of the endoderm and PGCs remains the same.

Finally, we asked whether PGCs mature over time. We measured the expression of several more mature PGC markers using qPCR and found that DPPA3 (stella) and DDX4 (vasa) show and increasing trend between 48h and 96h indicating that after their initial specification, PGC development continues (Fig. 2F). Another mature PGC marker, DAZL did not show significant expression, which is consistent with(Irie et al. 2015) which detected DAZL only in embryonic gonadal PGCs. In this context the increase of DDX4 is surprising since(Irie et al. 2015) also found DDX4 to only be expressed in gonadal PGCs and not in their hPGCLCs.

### PGCLCs share requirement for sustained BMP signaling with amnion-like cells

BMP and WNT signaling are known to be important for PGC specification, and the duration of exogenous WNT signaling is known to influence the decision between PGCLC versus primitive streak-like fates(Kobayashi et al. 2017; Sasaki et al. 2015), but the specific timing of the interplay between these two pathways is not well understood. We asked whether BMP and WNT act primarily through direct activation of PGC genes, or indirectly through induction of secondary signals, and whether the timing and duration of these signals matters.

First, we inhibited BMP signaling after 24h with the BMP receptor inhibitor (BMPRi) LDN193189 (Fig 3A-E, SI Fig. 5A-F). This led to a loss of PRDM1, a reduction in TFAP2C and outward displacement of TBXT. Notably, it also gave rise to a new FOXA2+SOX17-population at 42h (Fig. 3DE, SI Fig 5EF), which given the developmental stage and co-expression of TBXT may be axial mesoderm.

**Figure 3:**
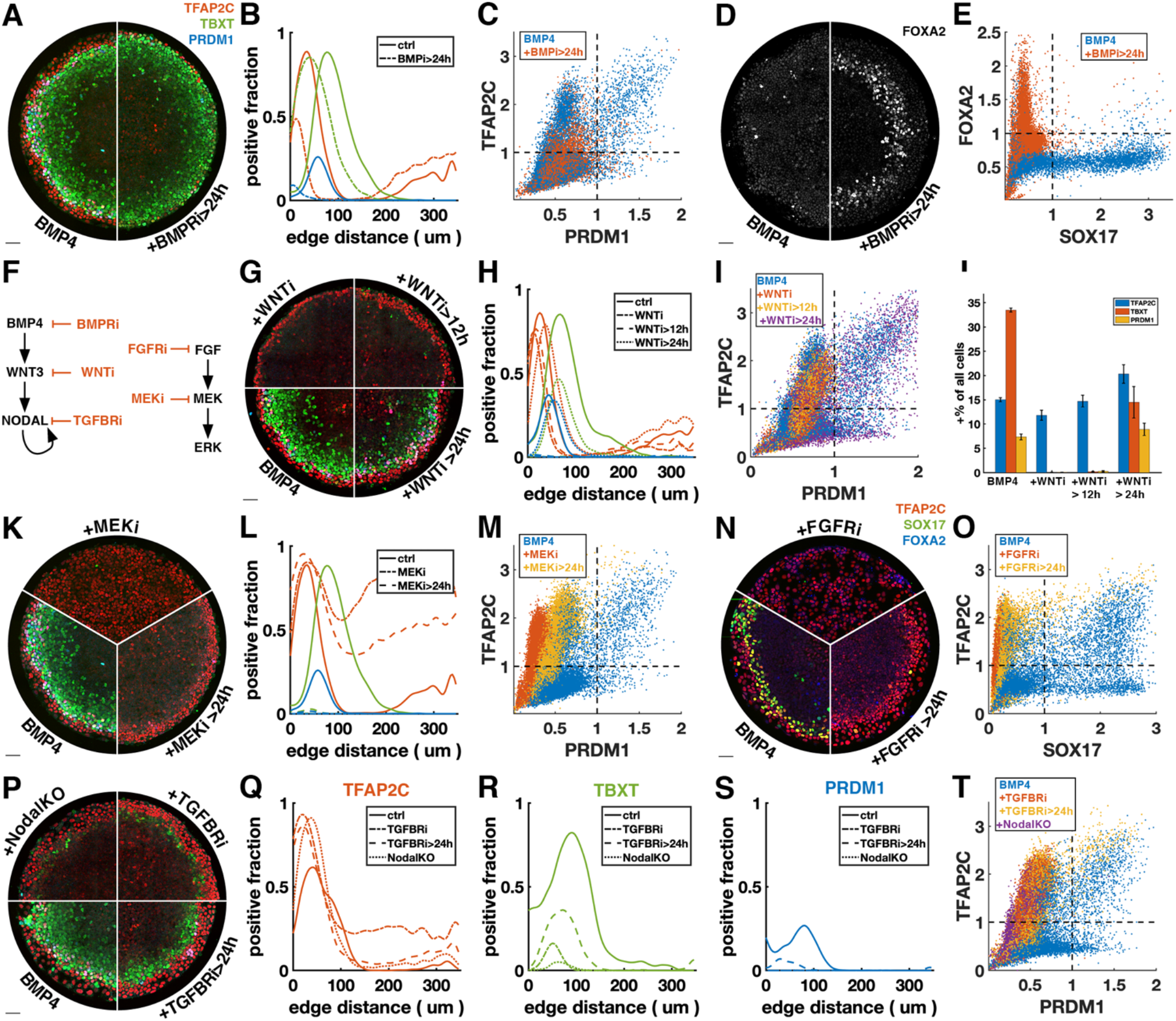
PGCLCs require sustained BMP, NODAL, and FGF but only brief WNT signaling. Each row shows staining and quantification of PGC markers after perturbation of different pathways. Error bands in spatial distributions are omitted for clarity but are similar in magnitude to Fig. 1c,e. **a-e)** BMP4-treated colonies with or without BMP-receptor inhibition after 24h (BMPRi, LDN-193189 250nM) shows loss of PGCLCs (a-c) and emergence of FOXA2+SOX17-population. **f)** Diagram of signaling hierarchy (black) and perturbations in this figure (red). **g-j)** WNT inhibition using iwp2 5uM after 0, 12 and 24h showing PGCLC specification only requires WNT signaling between 12 and 24h. **k-o)** Inhibition of FGFR (PD-173074 1uM) or MEK (PD-0325901 5uM) at 0 and 24h showing complete loss of PGCLCs in both cases. **p-t)** Inhibition of NODAL receptors (TGFBRi, SB-431542 10uM) at 0 and 24h and NODAL knockout (NODALKO) showing complete loss or severe reduction of PGCLCs. Images of each channel separately are shown in SI Fig. 5. Scalebar 50um.

Many of the effects of BMP4 are known to be indirect through the BMP-WNT-NODAL cascade. To test if the effect of LDN on PGC formation is direct or indirect, we next blocked WNT production (WNTi) downstream of BMP by IWP2 at different times (Fig. 3F-J, SI Fig. 5G). Surprisingly, blocking WNT production after 24h has almost no effect on PGC production, even though, as previously described, primitive streak markers are significantly reduced(Chhabra et al. 2019). This suggests that endogenous WNT signaling after 24h primarily serves primitive streak differentiation and the effect of BMPRi on PGCLC specification after 24h is the direct impact of BMP. PGCLCs share this dependence on sustained BMP signaling with AmLC, which were previously shown to require continuous BMP signaling past 24h(Chhabra et al. 2019; Nemashkalo et al. 2017).

As expected, WNT inhibition for the full duration of the experiment completely eliminates expression of TBXT and PRDM1. Strikingly, blocking WNT signaling after 12 hours has the same effect as blocking it at 0h, indicating that the role of WNT signaling in PGC specification is restricted to the window between 12 and 24 hours after addition of BMP4 (Fig 3G-J). We also noticed a reduction in the width of the TFAP2C expressing outer ring in these conditions (Fig. 3GH). However, quantification showed that the number of TFAP2C positive cells was not reduced, implying that the cells were more densely packed (Fig 3J). This suggests the change to a more spread-out morphology that is typically observed in the outer cells depends directly or indirectly on WNT signaling.

### NODAL and FGF signaling are required on the second day of PGCLC specification

Directed differentiation protocols for PGC typically consist of a brief period of exposure to primitive streak inducing signals WNT and NODAL to induce the incipient mesoderm like state (iMeLC) followed by BMP. Therefore, we asked whether the other signals that are required to specify mesoderm, FGF and NODAL, are also only required during the first 24h to induce PGCs.

First, we inhibited FGF signaling with FGF receptor inhibitor (FGFRi) PD173074 and MEK signaling (MEKi) with PD0325901after 0h and 24h (Fig. 3K-O, SI Fig H-I). The effects were very similar suggesting FGF specifies cell fate primarily through the MEK/ERK pathway and that the MEK/ERK pathway is primarily activated by FGF. When inhibiting FGF/ERK at 0h, TFAP2C expression becomes uniform throughout the colony and SOX17 expression is eliminated. When adding the inhibitor at 24h, TFAP2C expands inward and goes up in the colony center but is not uniform and a small number SOX17+TFAP2C+ PGCLCs is present. Our observation of inward expansion of TFAP2C is consistent with findings in zebrafish, where TFAP2C was found to be a direct target of BMP whose expression is excluded from the margin by FGF/ERK signaling(Rogers et al. 2020), suggesting conserved regulation by signaling of this gene despite diverging functions.

Next, we inhibited NODAL signaling with TGF-beta receptor inhibitor (TFGBRi) SB431542 (Fig 3. P-T, SI Fig 5J). NODAL inhibition for the full duration of the experiment eliminates PRDM1 completely and largely eliminates TBXT, while TFAP2C goes up inside the colony, similar to FGF inhibition, suggesting that either FGF and NODAL both restrict BMP response to the edge, or that one of these signals modulates the other. Inhibition at 24h severely reduces PRDM1 and TBXT expression but leaves a clearly defined ring of TBXT expression. This indicates that while WNT signaling is only required during the first 24h, both NODAL and FGF are also required at early and later times. Because there is TGF-beta in the differentiation media, TGFBRi not only blocks endogenous NODAL, but also the exogenous TGF-beta. To distinguish these effects, we looked at NODAL-/- cells (Fig. 3P-T) (Chhabra et al. 2019). While the panel of markers appeared similar to TGFBRi treated WT cells, TFAP2C expression in the center did not go up as much, indicating that in the absence of endogenous NODAL, low doses of TGFb in the differentiation media suppress TFAP2C and possibly BMP response more generally in the colony center.

### Exogenous Activin rescues PGCs in the absence of endogenous WNT or NODAL in a dose- and time-dependent manner

Given that WNT induces NODAL, it is possible that the main role of WNT for PGCLC specification is to induce NODAL. While it is known that NODAL alone does not induce differentiation, it is unclear what happens in combination with BMP. We therefore tested if we could rescue the loss of PGCs in WNTi by adding Activin to exogenously stimulate the NODAL pathway. PGC induction was indeed partially rescued by intermediate doses of Activin with robust expression of SOX17 and TFAP2C in a slightly reduced number of cells relative to BMP4 only (Fig 4A-B). This is surprising considering the literature that has emphasized the role of WNT signaling in hPGCLC induction(Hancock et al. 2021; Kobayashi et al. 2017). It was recently suggested that the main role of WNT is to induce EOMES (Yoney et al. 2021; Kojima et al. 2017). To test if it is possible to induce EOMES in the absence of WNT, we treated hPSCs with Activin, BMP4, and WNT3a for 4h in the absence and presence of inhibitors of the other pathways. We found that EOMES expression was also induced by BMP4 and Activin, although less strongly than by WNT3a, which may explain the presence but reduced numbers of PGCLCs (Fig. 4C). Although we cannot rule out that some WNT signaling still occurs in the presence of WNTi, the complete loss of PGCs and PS markers with WNTi in the absence of Activin, and significant rescue in the presence of Activin still suggests a key role for NODAL signaling.

**Figure 4:**
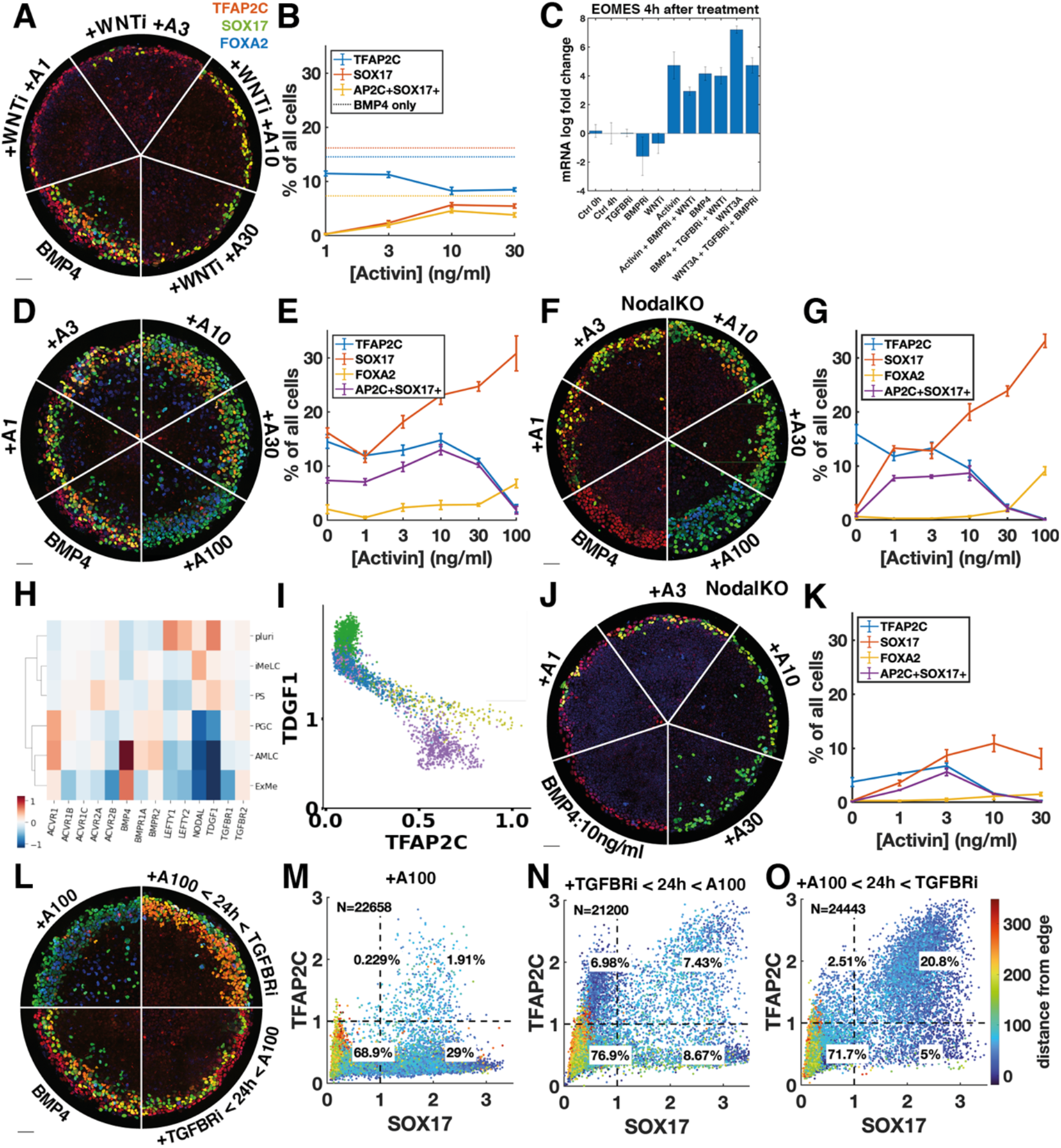
Exogenous Activin rescues PGCs in the absence of endogenous WNT or NODAL in a dose- and time-dependent manner. **a-b)** WNT inhibition (WNTi, iwp2) with different doses of Activin, e.g. A3 = 3ng/ml Activin. **c)** qPCR for EOMES relative to GAPDH, 4h after stimulation of pluripotent cells with different morphogens and inhibitors. Activin 50n/ml, BMP4 50ng/ml, WNT3a 300ng/ml, for notation also see methods and Fig. 3. **d-g)** Effect of treatment with Activin in WT and NODALKO cells on expression of PGC and endoderm markers. **h)** differential expression from scRNA-seq for NODAL and BMP receptors, as well as NODAL, BMP, and Lefty. **i)** scatterplot of TDGF1 vs. TFAP2C with colors corresponding to clusters in Fig. 1J. **j-k)** Effect of Activin treatment on NODALKO cells combined with a lower dose of BMP4 (10ng/ml vs. 50ng/ml in all other figures). **l-o)** Effect of 100ng/ml Activin for 42h, only during the first 24h, or only after 24h. Images of each channel separately are shown in SI Fig. 6. Scalebar 50um.

We then asked whether NODAL signaling might generally be the limiting factor in directing TFAP2C positive cells to PGCLC fate and treated colonies with different doses of Activin in addition to BMP4 (Fig 4DE). We found a moderate increase in PGCLCs at intermediate doses of Activin while at high doses TFAP2C was lost and SOX17+FOXA2+ cells appeared instead. NODAL autoactivates, so it is not clear how endogenous NODAL downstream of Activin changes and contributes to its effect. We therefore repeated this experiment in NODALKO cells (Fig. 4FG). We found that low doses of Activin rescue PGCLCs with numbers similar to wild type BMP4 treated cells, while higher doses behave very similar to wild type cells treated with the same dose of Activin. The similarity between NODALKO and wild type cells suggests feedback reduces the additive effect that might have been expected from Activin plus endogenous NODAL. To identify possible candidates for this feedback we looked at differential expression of NODAL, BMP and their receptors and inhibitors in the scRNA-seq data and found severely reduced expression of the NODAL co-receptor TDGF1 in the AmLC and PGCLC clusters (Fig. 4HI). This would desensitize those cells to NODAL but still allow strong response to Activin, which does not require TDGF1. We also noticed a striking upregulation of BMP-specific type 1 and 2 receptors ACVR1, BMPR1A and BMPR2, suggesting increased sensitivity to BMP4.

The dose-dependent effect of Activin treatment may be absolute if there are gene activation thresholds related to e.g., binding affinities of Smad2, or it may be relative to BMP, with BMP and NODAL signaling competing to activate and suppress TFAP2C. To test this, we reduced the dose of BMP4 from 50ng/ml to 10ng/ml and repeated the experiment (Fig 4JK). The lower dose of BMP4 induced less TFAP2C expression which limited PGC induction. Furthermore, much lower doses of Activin were sufficient to eliminate TFAP2C expression, indicating that the relative signaling levels determine the fate.

Unlike endogenous NODAL, high enough exogenous Activin eliminates TFAP2C and induces FOXA2 by 42h. We asked whether this is due to increased signaling levels or whether it is due to changes in timing and duration. Endogenous NODAL is activated with a delay downstream of BMP and WNT and does not reach high levels until after 24h(Heemskerk et al. 2019; Chhabra et al. 2019). At that point it is possible that the cells with the highest level of BMP signaling on the outside of the colony are already committed to AmLC fate and can no longer respond to NODAL or be converted to PGCLCs. Similarly, FOXA2 expression may require longer duration NODAL signaling that is not achieved by 42h if signaling starts at 24h. To test this, we treated cells with TGFBRi for the first 24h followed by a high dose of Activin in the last 24h (Fig. 4L-N). Consistent with our hypothesis, and in contrast to Activin treatment for the duration of the experiment, this did not eliminate TFAP2C or induce FOXA2 but instead induced PGCLCs in numbers similar to the BMP4-only control.

Combined, this data suggested that PGC specification requires that NODAL signaling should be activated in cells expressing TFAP2C to induce SOX17 before they commit to AmLC fate, but that prolonged high level NODAL signaling suppresses TFAP2C and activates FOXA2 to give rise to endoderm. Therefore, we predicted that a high dose of activin during only the first 24h would be able to convert all TFAP2C positive cells to PGCs without inducing endoderm. Indeed, we were able to double the fraction of PGCs to about 20% by 24h of Activin exposure with Activin/NODAL signaling inhibited after removing Activin (Fig. 4LO). As before, we repeated this experiment with NODALKO cells and obtained similar results to WT (SI Fig 6F-H).

In summary, we provide evidence that an important part of the role of WNT in PGCLC induction is indirect by inducing NODAL, and that the balance and relative timing of the NODAL and BMP pathways plays a crucial role in inducing TFAP2C and SOX17 to specify PGCs.

### Control of colony size dramatically improves PGCLC differentiation efficiency

As is clear from Fig. 4L-O, even with high doses of BMP and Activin expression of TFAP2C and SOX17 are confined to a 100um-wide ring or less on the edge of the colony that accounts for at most 30% of the cells. Response to exogenous BMP and Activin are well understood as an ‘edge effect’, excluded from the center by receptor localization and inhibitor production(Etoc et al. 2016). Therefore, we hypothesized that by reducing colony size thereby enhancing the total proportion of cells within a colony that are in contact with the edge, that we would be able to induce higher proportions of cells to express SOX17 or TFAP2C and get much larger fractions of PGCLC induction. To test this, we differentiated colonies ranging from 300um to 100um in diameter (Fig. 5A-C). We observed increasing fractions of PGCLCs as the diameter decreases, reaching a maximum for 100um colonies at about 50% SOX17+TFAP2C+ (Fig.5D) or TFAP2C+PRDM1+NANOG+ (Fig. 5EF). Moreover, PS-like cells as marked by high EOMES were eliminated in 100um colonies and nearly all cells expressed TFAP2C, suggesting that the non-PGCLCs are AmLCs. Since complex current protocols yield 20-30% PGCLCs, it is surprising that BMP4 treatment combined with controlled colony geometry alone would yield 50% PGCLCs. We asked if the yield would increase further by pre-differentiating cells to iMeLC with 12h of WNT and NODAL activation as is done in current protocols, or by treating with Activin for the first 24h as we did in Fig. 4. Indeed, the fraction of PGCLCs increased to 70% by pre-differentiation (Fig. 5G). Surprisingly, treatment with Activin did not have the same effect as for large colonies and left the number of PGCLCs unaffected (Fig. 5H). In summary, combining current protocols with geometric control using micropatterning more than doubles their efficiency.

**Figure 5:**
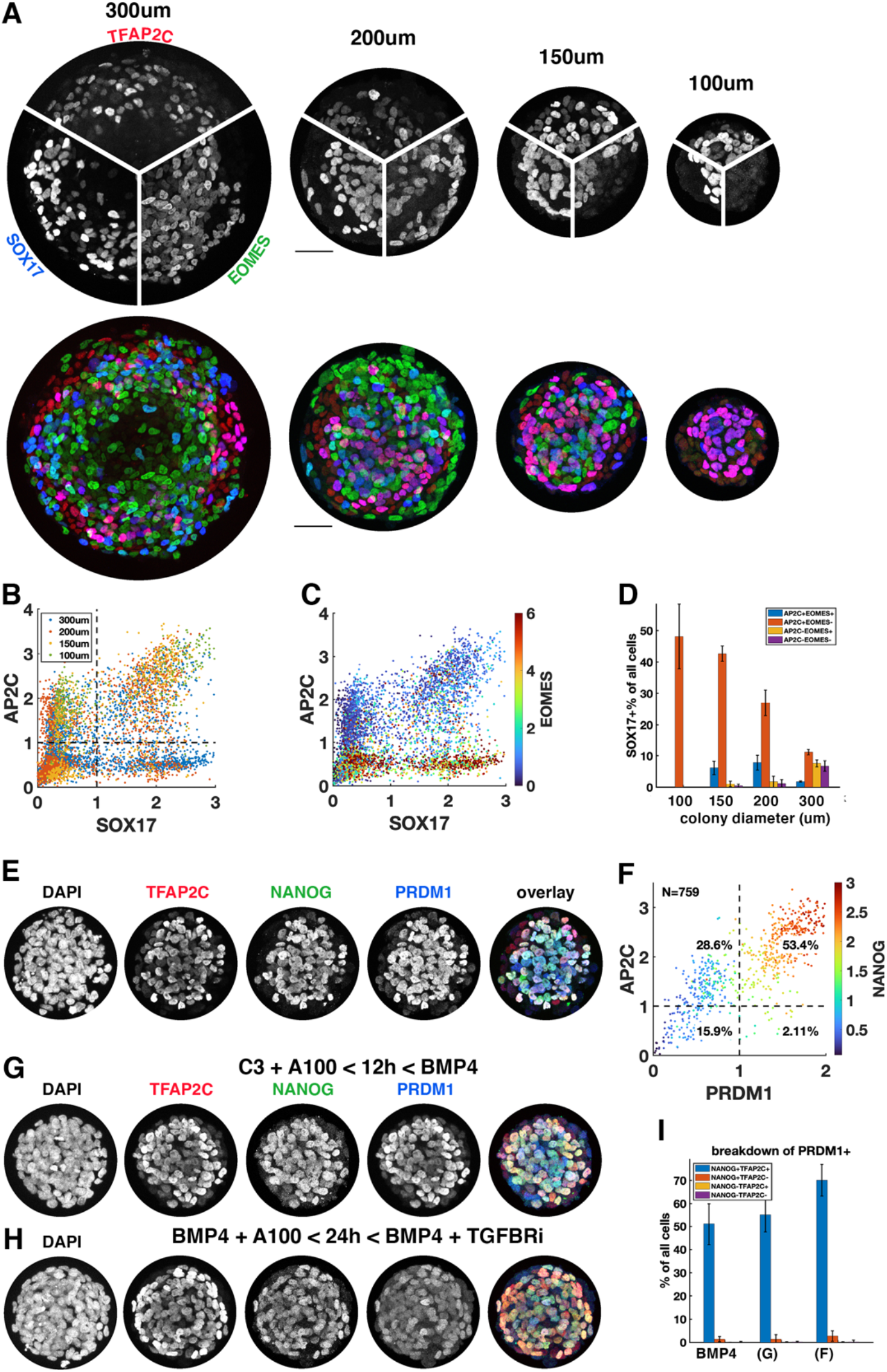
Control of colony size dramatically impacts the fraction of PGCLCs. **a-d)** Different diameter colonies stained for TFAP2C, SOX17, EOMES at 42h and quantification, **b)** SOX17 vs. TFAP2C scatter plot colored for colony size **c)** same plot colored for EOMES expression. **d)** SOX17+ subpopulations for each colony diameter. **e-f)** 100um colony stained for TFAP2C, NANOG, PRDM1 at 48h and quantification (17 colonies per condition). **g-i)** 100um colonies differentiated with alternative protocols and quantification. C3 = 3uM CHIR-99021, other notation is like in figure 4. Scalebar 50um.

### A network of cross-repressive cell fates driven by BMP and NODAL signaling explains perturbations of PGCLC induction

Our experiments suggest that cells interpret the ratio of BMP and NODAL signaling levels as well as their relative timing and duration through a network of mutually repressive fates that are acquired in a switch-like manner. Although WNT is required in combination with NODAL to induce PS-like fates, for PGCLC specification its primary role is to induce NODAL, and most of our results can be explained without considering WNT. Moreover, differentiation of AmLCs on the colony edge does not require NODAL, and PS-like fates do not (directly) require BMP, while PGCLCs positioned in between require combined induction of both BMP and Nodal target genes, in particular TFAP2C and SOX17.

Intuitively, this would explain the unperturbed wild type pattern as follows. First, higher BMP signaling on the colony edge induces TFAP2C and amnion genes faster than further inside; then, with a delay that depends on distance from the edge, Nodal signaling turns on in all cells at similar levels and induces SOX17 and PS-like genes. Cells on the outside reach a threshold to stably switch on amnion-like genes and repress other fates before SOX17 and PS-like genes are significantly induced. Cells slightly further inside reach high enough levels of both TFAP2C and SOX17 to stably switch on PGCLC genes and repress other fates, while cells even further inside never significantly activate amnion genes or TFAP2C and after a longer exposure to Nodal will reach a threshold to commit to a PS-like fate. Combined induction of PS-like genes and SOX17 may specify endoderm. The effect of perturbations of Activin/NODAL timing and duration is then naturally explained: early treatment with Activin combined with BMP can induce high enough levels of both SOX17 and TFAP2C in cells on the colony edge to make them PGCLCs before they commit AmLC, but if the duration of Activin exposure is too long PS-like genes dominate and suppress PGCLC fate. On the other hand, Activin exposure in the second 24h coincides with the timing of endogenous NODAL and has little effect on fate decisions.

To test if this intuitive model holds up more rigorously and also explains our other perturbations of BMP and NODAL signaling, we identified a minimal mathematical model for the gene regulatory network specifying PGCs downstream of previously determined BMP, NODAL, and Activin signaling profiles. Previous work showed that after initial uniform activation, BMP signaling is restricted to a stable gradient from the colony edge and that a region of high endogenous Nodal signaling expands into the colony from the edge at constant velocity starting around 24h (Etoc, Heemskerk et al. 2019, Fig. 6A). Like BMP, the response to exogenous Activin forms a gradient from the edge. For simplicity we did not include WNT in our model because our observations appear to at least be qualitatively explained without WNT. We also did not include FGF and its perturbations since we lack data on the spatiotemporal profile of FGF signaling. Moreover, we modeled PGCLCs as TFAP2C+SOX17+ and treated SOX17 and PRDM1 as interchangeable in terms of the observations explained by the model, knowing that PRDM1 is downstream of SOX17. For Activin treatment we chose to model NODALKO cells since the combined Nodal and Activin signaling is unknown and experiments suggest a feedback that results in only minor differences between WT and NODALKO. Specific choices made in the construction of the model are further detailed in the supplementary text.

**Figure 6:**
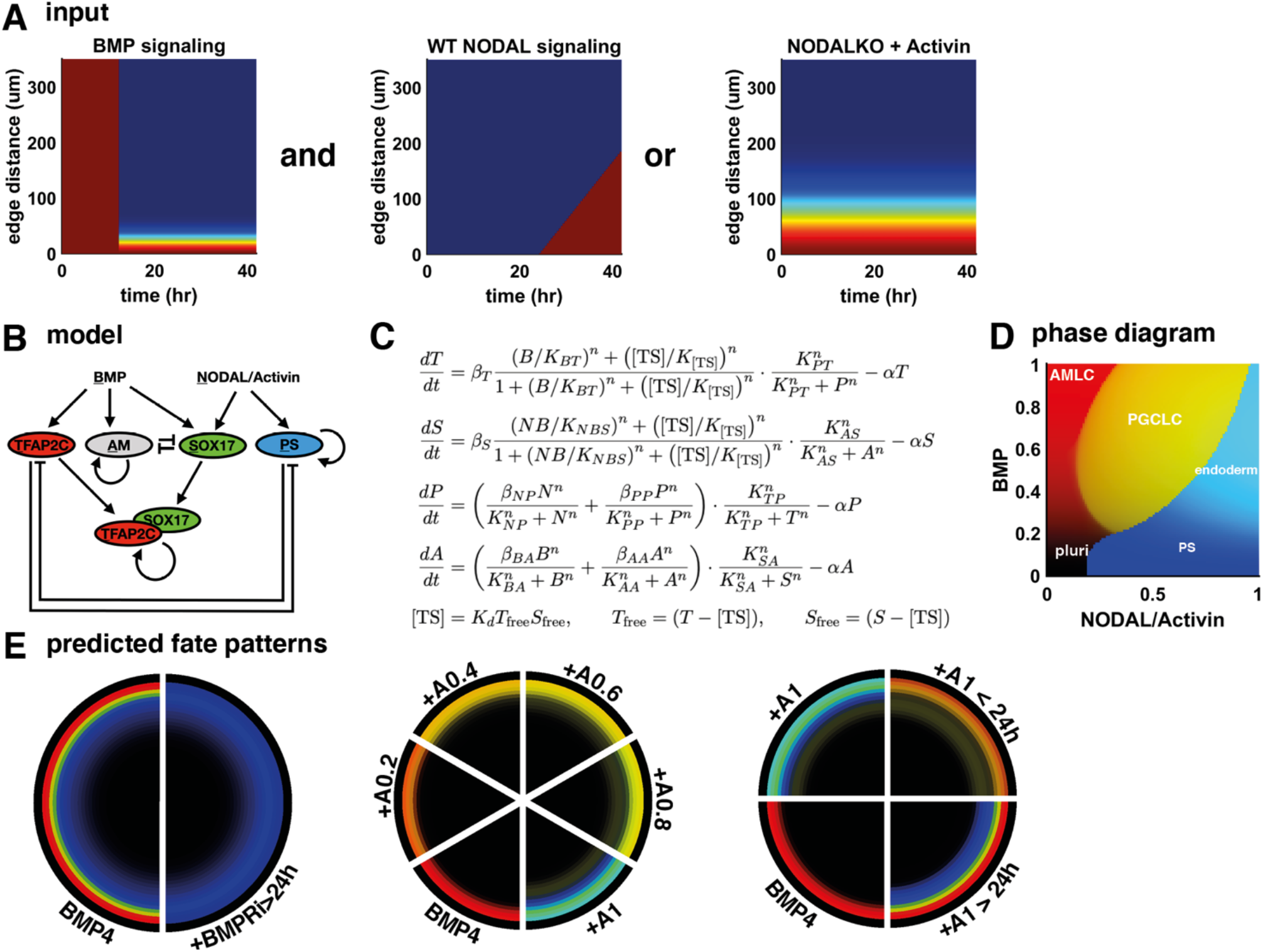
A network of cross-repressive cell fates qualitatively explains NODAL perturbations. **a)** Input signaling profile in space and time. **b)** Diagram of the model. **c)** Definition of the model. **d)** Phase diagram showing predicted expression of cell fate markers at 42h for different levels of BMP and NODAL activation. **e)** Cell fate patterns predicted by the model from the input signaling profiles with different perturbations. Compare with data in Fig. 3A, 4F, and SI Fig. 6F.

The structure of the model is shown in Fig. 6BC, and the cell fate markers induced by constant levels of NODAL or BMP signaling are shown in the phase diagram in Fig. 6D. We then tested if this model explains the effect of our signaling perturbations on cell fate patterns and found qualitative agreement (Fig 6E). We also tested various simpler models, for example lacking the competition between PGCLCs and the neighboring AmLC and PS-like fates and found that these do not explain the data. In summary, we find that the mathematical model strongly supports our interpretation of the data.

## DISCUSSION

In this study we have shown that PGCs form in a very reproducible manner in BMP4-treated micropatterned hPSCs and are part of a stereotypic spatial organization that positions them in between extraembryonic cells that may be amnion like and cells expressing primitive streak markers, similar to their location *in vivo*. Because they are close to the colony edge we used micropatterning to create small colonies and were able to get much greater fractions of PGCLC differentiation than previously described. This potentially also explains the advantage in using ROCK inhibitor in PGCLC differentiation noted by some groups(Sebastiano et al. 2021) since that keeps cells from forming densely packed colonies and makes the majority of cells behave like the colony edge. However, micropatterning provides a more tightly controlled way to achieve this effect. By incorporating 12 hours of iMeLC differentiation we were able to achieve 70% efficiency compared to 10-30% described in the literature. It is possible that further protocol optimization of micropatterned differentiation could further increase the yield.

In a major advance, a microfluidic system for amniotic sac development was recently shown to give rise to hPGCLCs and was used to study the transcriptome of hPGCs(Zheng et al. 2019; Chen et al. 2019; Yang et al. 2021). However, quantitative studies of the signaling dynamics underlying specification of cell populations have mostly been performed in micropatterned hPSCs due to the simplicity of the system(Heemskerk 2019). The cells form a quasi-two-dimensional system, thereby creating optimal conditions for quantitative live cell microscopy and micropatterned substrates are easy to make and commercially available.

It was a surprise that in our hands, there was no clear definitive endoderm (DE) population present at 42h and the majority of SOX17+ cells that were originally thought to be endoderm are PGCLCs. However, we found that endoderm is present at 48h and continues to increase until 72h. It is possible the SOX17+TFAP2C- cells at 42h will continue to differentiate to DE, which would require lineage tracing, but neither immunofluorescence nor scRNA-seq shows expression of DE markers like FOXA2 or HEX at 42h. When endoderm arises, it localizes close to the PGCs and is still in a location where high BMP is expected. Therefore, the puzzle of endoderm localization in micropatterned hPSC colonies is not fully resolved by our study. However, NODAL and BMP have not been measured after 48h, making it is possible that BMP signaling is excluded from this region at later times.

The fact that endoderm forms primarily between 48h and 72h and that we observed further maturation of PGCLCs suggest more broadly that pattern formation and differentiation continue during this time window which so far has not been studied. It will be interesting to also consider what happens with the mesodermal lineages during this time window.

A different study performed scRNA-seq at 44h(Minn et al. 2020) and found an endodermal and PGC population. This may capture the earliest endoderm formation we observed. It is also possible that there are subtle differences in timing or differentiation potential that depend on the cell line and our data does show some variation between cell lines (SI Fig. 3). Differences may also be due to details of the protocol. For example, we have observed that the progression of marker expression can differ by a few hours depending on factors like the total media volume or small differences in the initial cell density. Future work on micropatterned stem cells could benefit from developing a consensus staging, where e.g., the first appearance of a TBXT ring, and the first expression of FOXA2, could define stages.

Our earlier work showed that endogenous NODAL does not form static gradients but moves into the colony like a wavefront with constant velocity, arguing against the classic model of pattern formation by concentration thresholds. Moreover, we found that response to Activin and NODAL is adaptive and that gene response depends in part on signal rate of change. Here we have observed both duration-dependent and dose-dependent PGCLC specification by Activin and NODAL. These findings are all consistent with our model in which gene expression depends on integrated signaling activity that could increase either through concentration, rate of concentration change, or duration. The integrated signaling over time is then interpreted by a gene regulatory network (GRN) to make cell fate decisions.

Although to our knowledge the interactions in our model for the GRN are consistent with the literature, there are several variations possible at the level of the model, and several molecular mechanisms that could be responsible for the behavior of the same model (SI text). For example, instead of directly, BMP could activate SOX17 indirectly through TFAP2C as suggested by(Chen et al. 2019). Future work will be able to refine this model to make more accurate predictions for the markers of interest and move towards quantitative predictions of PGCLC specification.

While the focus in hPGCLC differentiation has typically been on BMP and WNT, we showed that a major part of the role of WNT is to induce NODAL and that the effect of WNT inhibition on hPGCLC specification can be partially rescued by exogenous Activin. This further highlights the complex feedbacks between the paracrine signaling pathways that make it hard to directly interpret the effect of a signaling perturbation, and therefore the need for a quantitative approach. By establishing a highly efficient and reproducible differentiation platform and revealing how timing, duration, and dose of Activin/NODAL signaling affect hPGCLC specification, we have laid the foundation for future quantitative investigations of the interplay between different signaling pathways during PGCLC induction, and the downstream gene regulatory that interprets these signals to determine fate.

## METHODS

### Replicates

All experiments were performed at least twice, quantification was done on four or more colonies per condition. Error bars represent standard deviation over colonies unless otherwise stated.

### Cell lines

The cell lines used were the embryonic stem cell line ESI017 (XX), and the induced pluripotent stem cell lines PGP1 (XY), WTC11 (XY), MR30 (XX). The identity of these cells as pluripotent stem cells was confirmed by staining of pluripotency markers OCT3/4, SOX2, NANOG. All cells were routinely tested for mycoplasma contamination, and negative results were recorded.

### Pluripotent stem cell culture and differentiation

Pluripotent stem cells were cultured in mTeSR1 media (StemCelll Technologies) on Cultrex (R&D Systems) coated tissue culture plates. Whole colony routine passaging was done using L7(Nie et al. 2014) and single cell suspension for seeding experiments with were generated using Accutase. For micropatterned colonies, we followed(Deglincerti et al. 2016) using the chemically defined medium mTeSR1. Unless stated otherwise, BMP4 treatment was done with 50ng/ml. All experiments were done in micropatterned 18-well Ibidi slides made using the protocol(Azioune et al. 2009). All colonies were 700um diameter unless stated otherwise. We used the following reagents to modify signaling during pattern formation:

### Imaging and image analysis

Imaging was done on an Andor Dragonfly / Leica DMI8 spinning disk confocal microscope with a 40x, NA 1.1 water objective. Nuclei were segmented in individual z-slices based on DAPI staining using two different machine learning approaches: Ilastik(Sommer et al. 2011) and Cellpose (Stringer et al. 2021). We found Cellpose is highly accurate with segmenting the nuclei it finds, but it sometimes misses lower contrast nuclei, whereas Ilastik can be easily trained to find all nuclei but more frequently has trouble separating neighboring nuclei. Therefore, we combined the two segmentations in each z-slice giving preference to Cellpose. To get a 3D segmentation, we next linked segmentations in different z-slices using using a linking algorithm formulated as a linear assignment problem that takes into account the overlap, relative position of centroids, and second derivatives of area and intensity with respect to z of nuclear masks in consecutive z-slices to decide whether they belong to the same nucleus or not (see SI Methods for details). Using the resulting segmentation, we extracted mean intensities for each of the stained markers in each nucleus. For further analysis, the single cell expression data obtained this way was log(1+x) transformed similar to what is common for scRNA-seq analysis, for several reasons including reduction of the effect of outliers on the analysis. We separated population by thresholds in each marker, which while not perfect performed better than more advanced clustering methods. To determine a threshold between cells expressing or not expressing a marker, we fitted a Gaussian mixture model to the expression data of each gene separately, which worked better than fitting it to the combined gene expression due to the clusters not being sufficiently Gaussian in two or three dimensions. The number of Gaussians was determined automatically using the Bayesian Information Criterion, and the positive cells were taken to be those belonging to the Gaussian with the highest mean. This agreed well with the threshold but in many cases required some manual fine tuning based on the scatterplots (thresholds shown in all scatter plots). The data was then rescaled so that log(1+x_thresh) = 1 for visualization in scatterplots. All code will be available on github.com/idse/PGCs.

### Immunostaining

Coverslips were rinsed with PBS, fixed for 20min in 4% paraformaldehyde, rinsed twice with PBS, and blocked for 30 min at room temperature with 3% donkey serum and 0.1% Triton X-100 in 1× PBS. After blocking, cells were incubated with primary antibodies at 4°C overnight (Table S1) followed by three washes in PBST (PBS with 0.1% Tween 20). They were then incubated with secondary antibodies and DAPI for 30 min at room temperature and washed twice in PBST at room temperature. Antibodies can be found in tables 2,3.

**Table 1.**
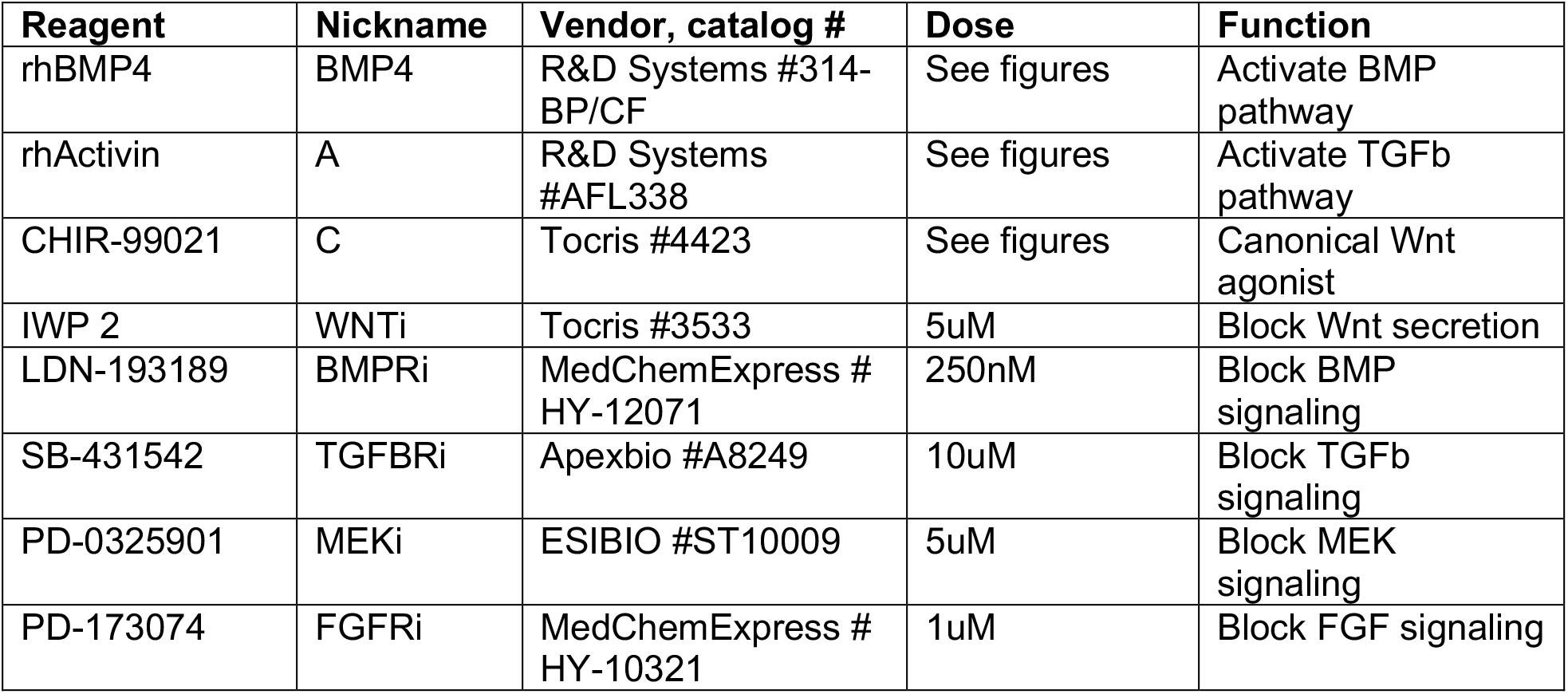
Cell signaling reagents.

**Table 2.**
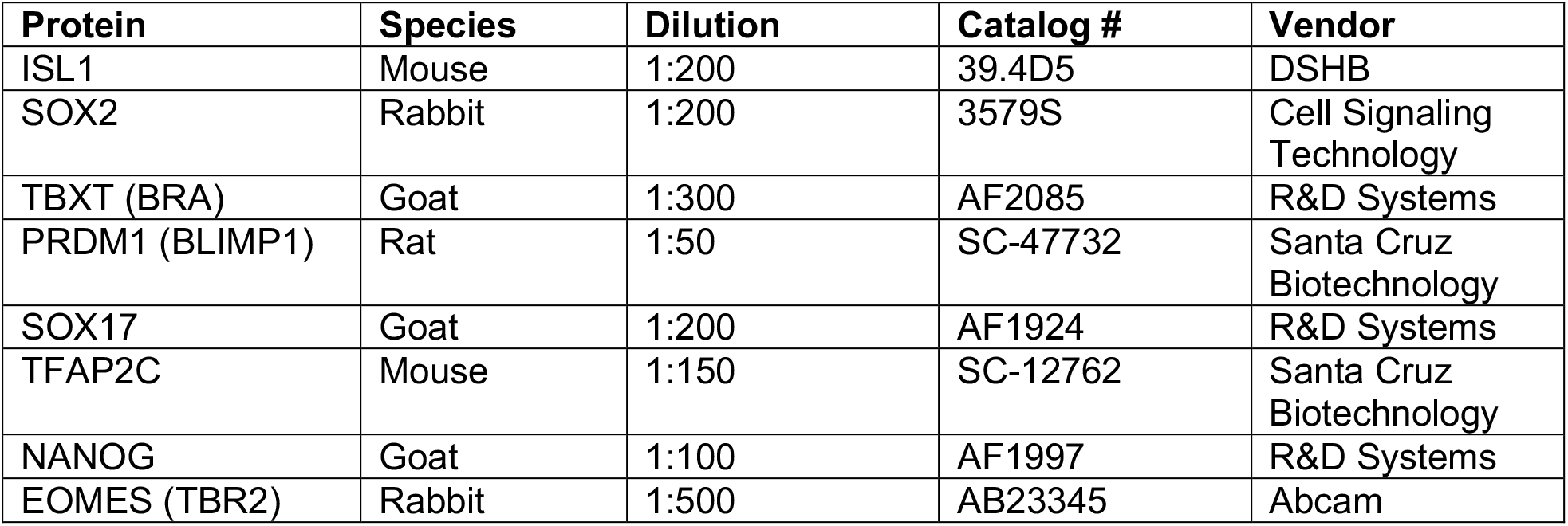
Primary antibodies used for immunostaining.

| Protein | Species | Dilution | Catalog # | Vendor |
| --- | --- | --- | --- | --- |
| ISL1 | Mouse | 1:200 | 39.4D5 | DSHB |
| SOX2 | Rabbit | 1:200 | 3579S | Cell Signaling Technology |
| TBXT (BRA) | Goat | 1:300 | AF2085 | R&D Systems |
| PRDM1 (BLIMP1) | Rat | 1:50 | SC-47732 | Santa Cruz Biotechnology |
| SOX17 | Goat | 1:200 | AF1924 | R&D Systems |
| TFAP2C | Mouse | 1:150 | SC-12762 | Santa Cruz Biotechnology |
| NANOG | Goat | 1:100 | AF1997 | R&D Systems |
| EOMES (TBR2) | Rabbit | 1:500 | AB23345 | Abcam |

**Table 3.** secondary antibodies

| Protein | Species | Dilution | Catalog # | Vendor |
| --- | --- | --- | --- | --- |
| Alexa Fluor 647 Anti-Goat | Donkey IgG | 1:500 | A21447 | Thermo Fisher Scientifc |
| Alexa Fluor 555 Anti-Goat | Donkey IgG | 1:500 | A21432 | Thermo Fisher Scientifc |
| Alexa Fluor 488 Anti-Mouse | Donkey IgG | 1:500 | A21202 | Thermo Fisher Scientifc |
| Alexa Fluor 647 Anti-Rat | Whole IgG | 1:500 | 112-605-167 | Jackson Immuno Research |
| Alexa Fluor 647 Anti-Rabbit | Donkey IgG | 1:500 | A31573 | Thermo Fisher Scientifc |
| Alexa Fluor 555 Anti-Rabbit | Donkey IgG | 1:500 | A31572 | Thermo Fisher Scientifc |

### qPCR

For qPCR experiments, ESI017 cells were grown in 24-well plates or 18-well ibidi slides. For Eomes response in Figure 4C, all treatments were done by taking part of the media from each well to dilute treatment reagents that were then added back to the well, in order to prevent effects from adding fresh media. RNA was extracted using Ambion RNAqueous-Micro Total RNA Isolation Kit and cDNA synthesis was performed with Invitrogen Super- Script Vilo cDNA Synthesis Kit according to the manufacturer’s instructions. Measurements were performed with SYBR green and the primers are in Table 4. GAPDH was used for normalization. In all cases, at least two biological replicates were performed and showed similar results. Error bars on qRT-PCR data are over technical triplicates from the representative biological sample set.

**Table 4.**
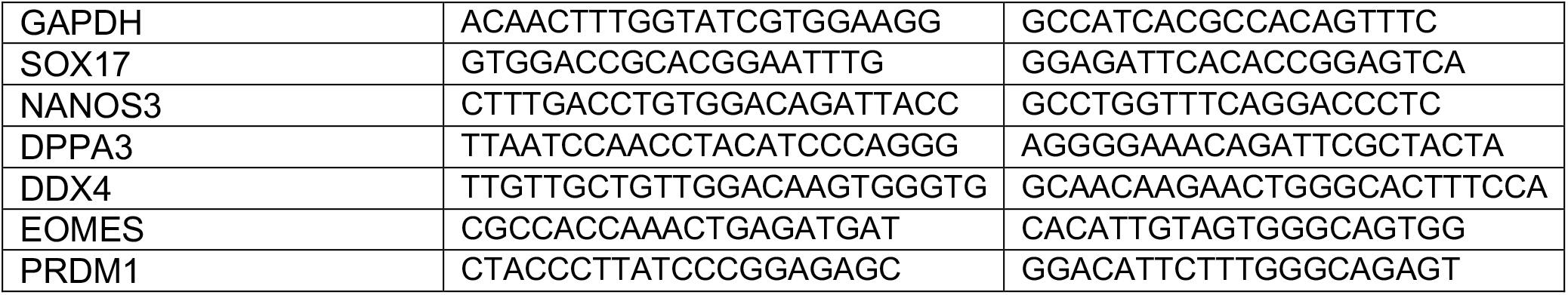
qPCR primers.

### scRNA-seq

Cells were collected using accutase and resuspended in ice cold PBS. Single-cell RNA-sequencing was performed by the University of Michigan Advanced Genomics Core. Cells were barcoded using the 10x Genomics Chromium system (part numbers 1000268, 1000120, 1000215). For quality control cDNA was quantified by Qubit High Sensitivity DNA assay and Agilent TapeStation. Sequencing was performed on the Illumina NovaSeq 6000 with NovaSeq S4 flowcell and Control Software version 1.7.0. Reads were aligned using cellranger-4.0.0 with the GRCh38 reference. Further processing was done in Python using the scprep package, the script used for all analysis which includes all parameters is included as a supplement. After filtering for library size to exclude empty droplets and duplets, 4254 cells were left. After excluding outliers for mitochondrial gene expression and excluding genes that were expressed in fewer than 50 cells, we were left with 4095 cells and 16151 genes. The data was then transformed using a sqrt which has a similar effect as the commonly used log(1+x) transformation without the arbitrary pseudocount to avoid singular behavior at zero. Rather than regress out various factors like cell cycle or pseudogenes that were not of interest or may confound analysis and lead to misinterpretation(Chhabra and Warmflash 2021), we made a list of 190 developmental genes of interest (SI table 1) that we used for visualization and clustering. We found both visualization and clustering to be more reliable and more informative this way and found our results to be very stable to adding genes or to removing genes from this list. Dimensional reduction for visualization was performed using PHATE, which preserves the global structure, i.e. lineage structure of the data better than UMAP without compromising local structure. For visualizing gene expression on PHATE plots and visualizing gene relationships using DREMI, we first performed denoising using MAGIC. Other analysis such as differential expression was performed on the full data. Data was scaled to zero mean and unit variance before performing differential expression analysis using Earth Mover’s Distance.

## Supporting information

Supplementary Text on Image Analysis

Supplementary Text on Mathematical Model

## ACKNOWLEDGEMENTS

We thank Aryeh Warmflash, Sue Hammoud, and Craig Johnson for discussions. NODALKO cells were a gift from Aryeh Warmflash. This work was supported by the Branco Weiss Fellowship – Society in Science, the University of Michigan Pioneer Postdoctoral Fellowship, the University of Michigan, and the National Institute of General Medical Sciences.

## SUPPLEMENTARY FIGURES

**Supplementary figure 1:**
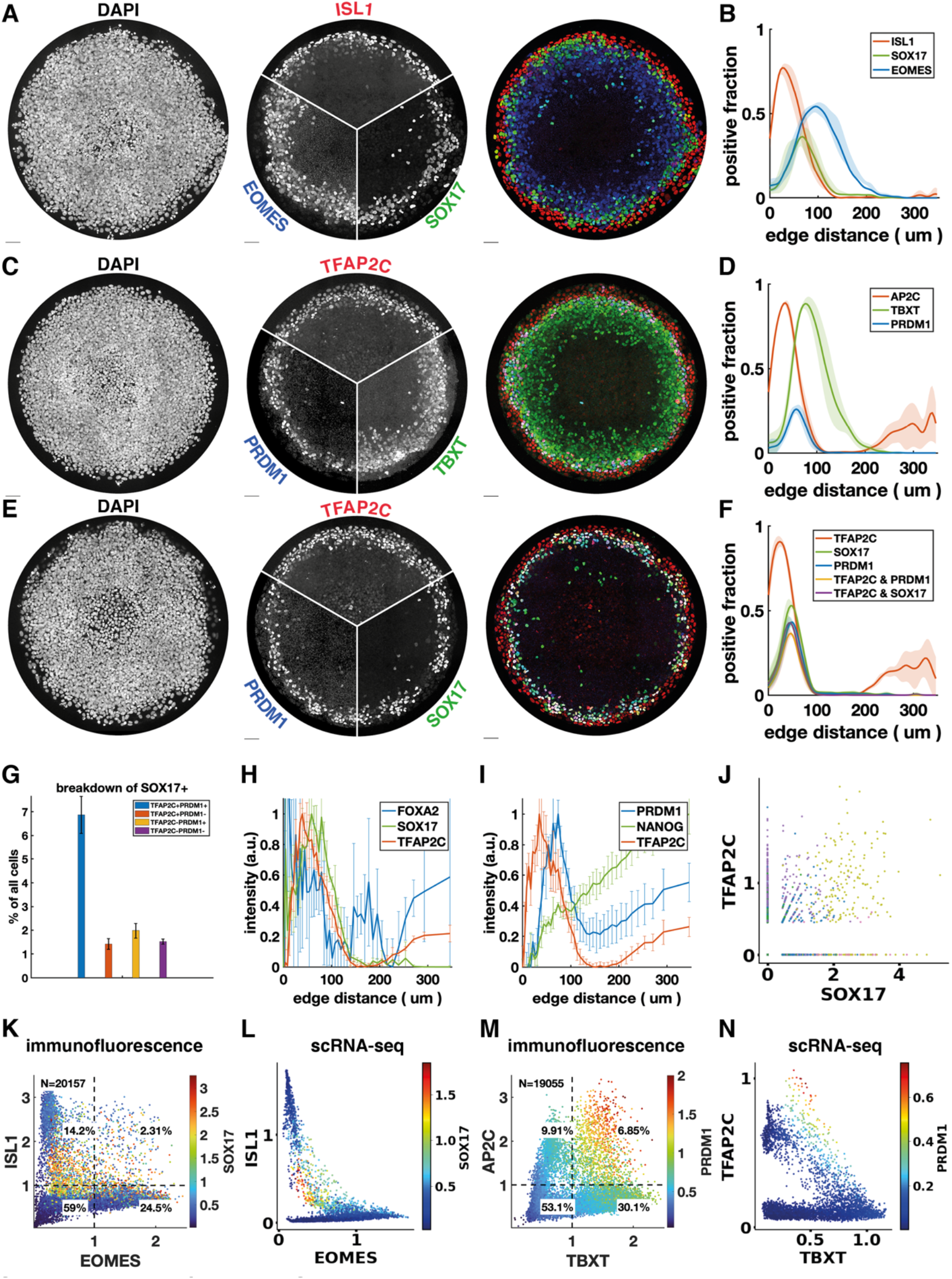
Quantitative relationships between marker genes with IF and scRNA-seq. **a-f)** Additional stainings of micropatterned colonies and spatial distribution of positive cells. **g)** breakdown of cell populations corresponding to Fig. 1g and SI Fig. 1ef. **h-i)** Normalized radial intensity profiles corresponding to Fig. 1ab. **j)** Scatterplot of TFAP2C vs. SOX17 from scRNA-seq data before denoising with magic, compare with Fig.1m. **k-l)** Scatterplots comparing the relationship between ISL1, EOMES, SOX17 measured with IF (k) and scRNA-seq (l). **m-n)** Scatterplots comparing the relationship between TFAP2C, TBXT, PRDM1 measured with IF (m) and scRNA-seq (n).

**Supplementary figure 2:**
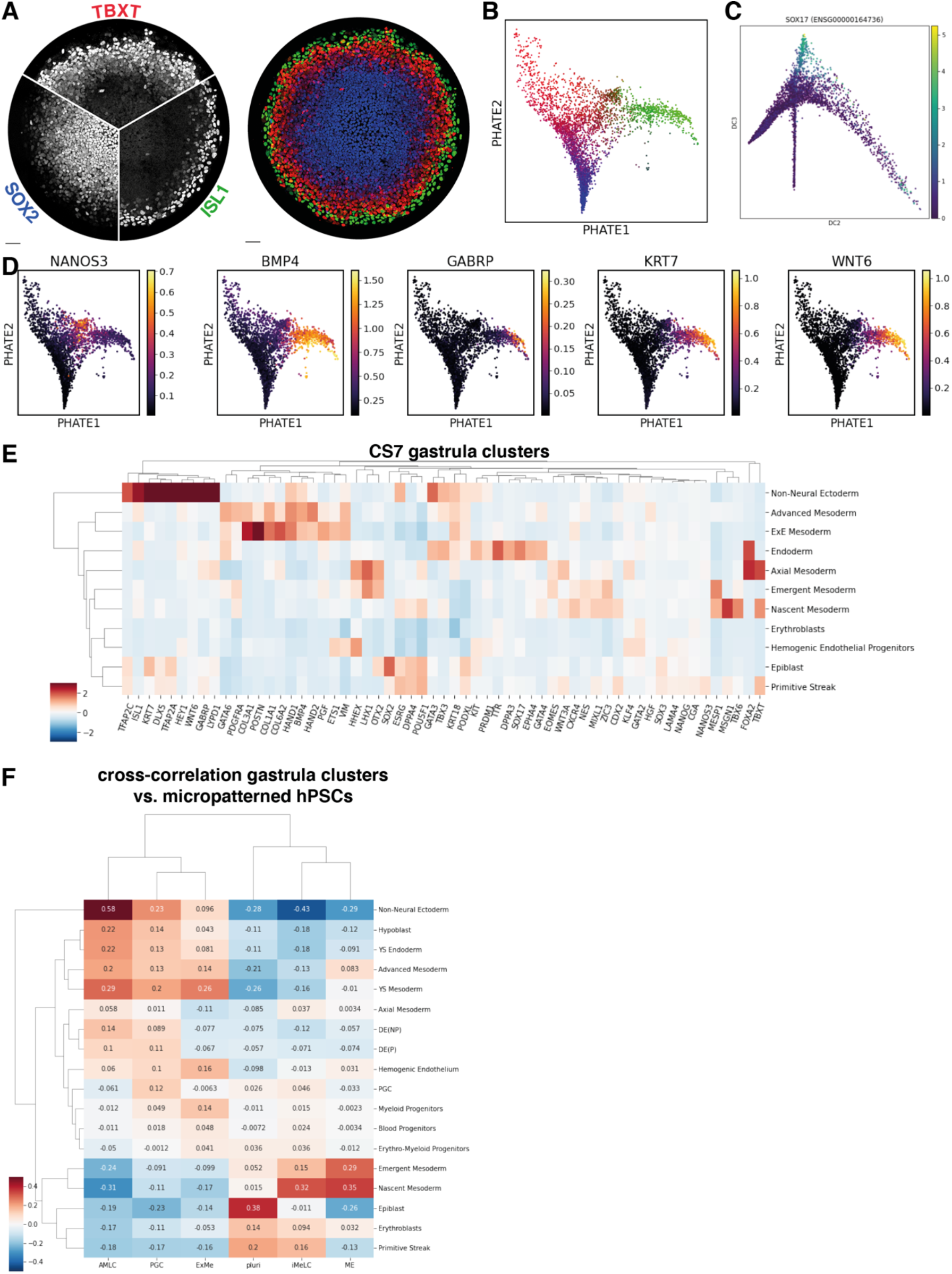
Additional scRNA-seq analysis, including correlation with CS7 human gastrula. **a-b)** comparison of IF for ISL1, TBXT, SOX2 with their expression on the PHATE visualization of the scRNA-seq data. **c)**PGCs budding off as a lineage along diffusion component 3. **d)** Expression of amnion and PGC markers on PHATE visualization. **e)** Differential expression of gastrula markers from Fig. 1k in clusters of CS7 human gastrula from Tyser et al. **f)** Correlation of highly variable genes between the clusters from micropatterned hPSCs and clusters from human gastrula.

**Supplementary figure 3:**
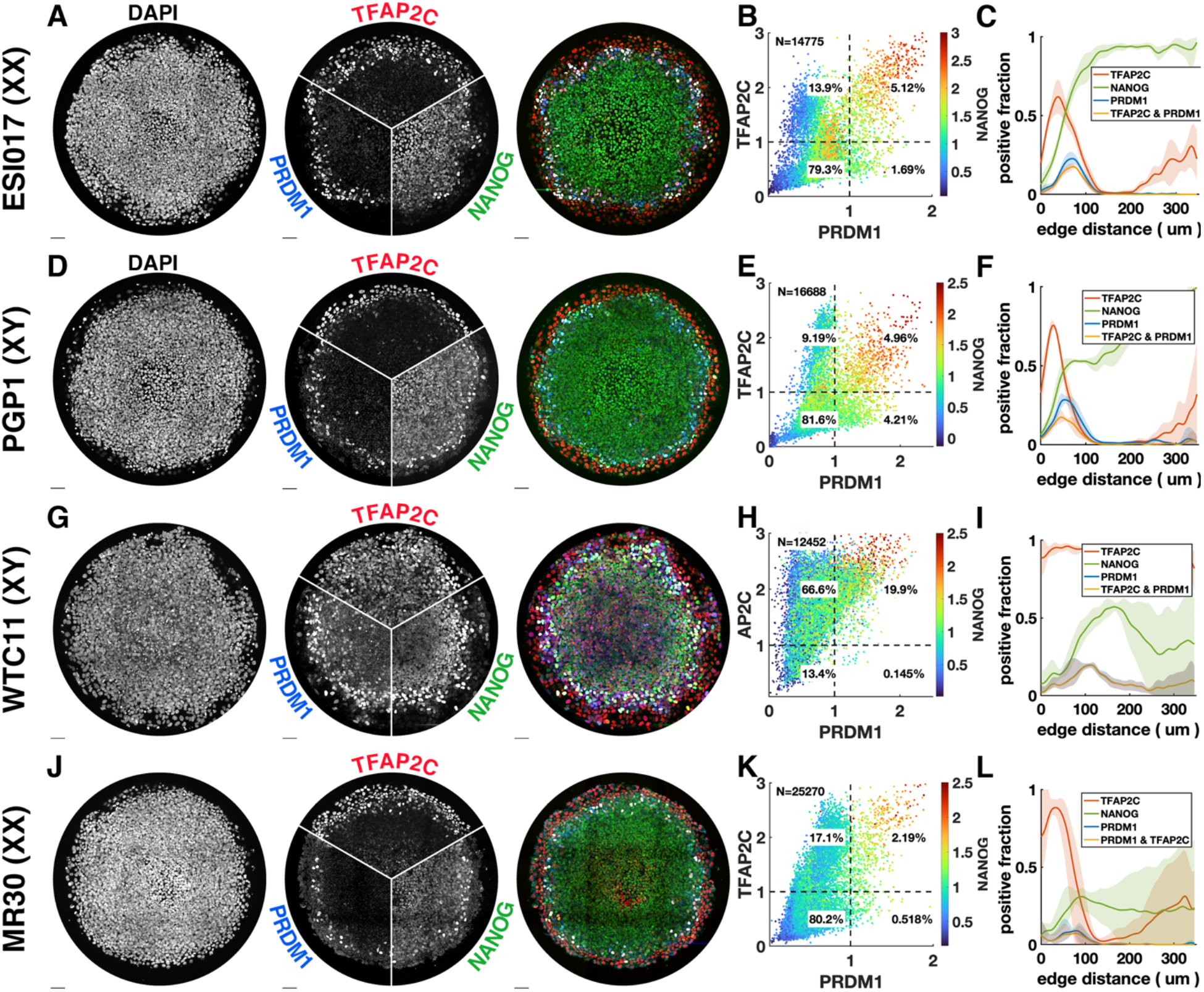
PGCLC differentiation in different male and female cell lines. Each row shows IF and quantification for PGC markers in different cell lines. **a-c)** ESI017, also shown in Fig. 1B, shown here for comparison. **d-f)** PGP1, male iPSC. **g-i)** WTC11, male iPSC. **j-l)** MR30, female iPSC. Strikingly, WTC11, a widely used iPSC line deviates from the norm. It consistently expresses TFAP2C throughout the colony and gives rise to higher numbers of PGCLCs.

**Supplementary figure 4:**
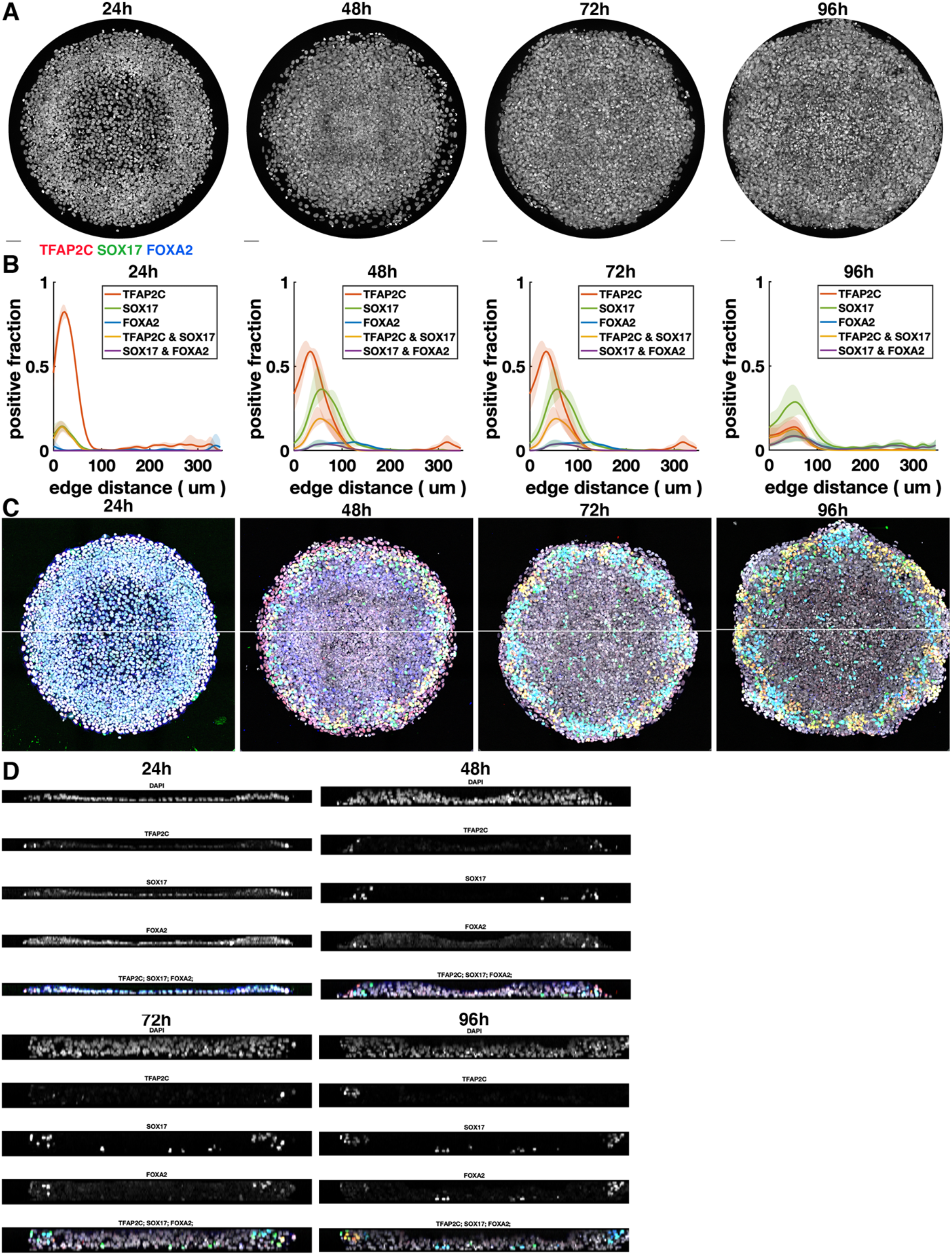
Additional images and quantification for time series up to 96h. **a)** DAPI staining for colonies shown in Fig. 2ab. **b)** Radial profile of marker expression at different times. **c)** overlay of colonies from Fig. 2ab including DAPI, showing location of cross-section in d. **d)** Cross sections through colonies at different times.

**Supplementary figure 5:**
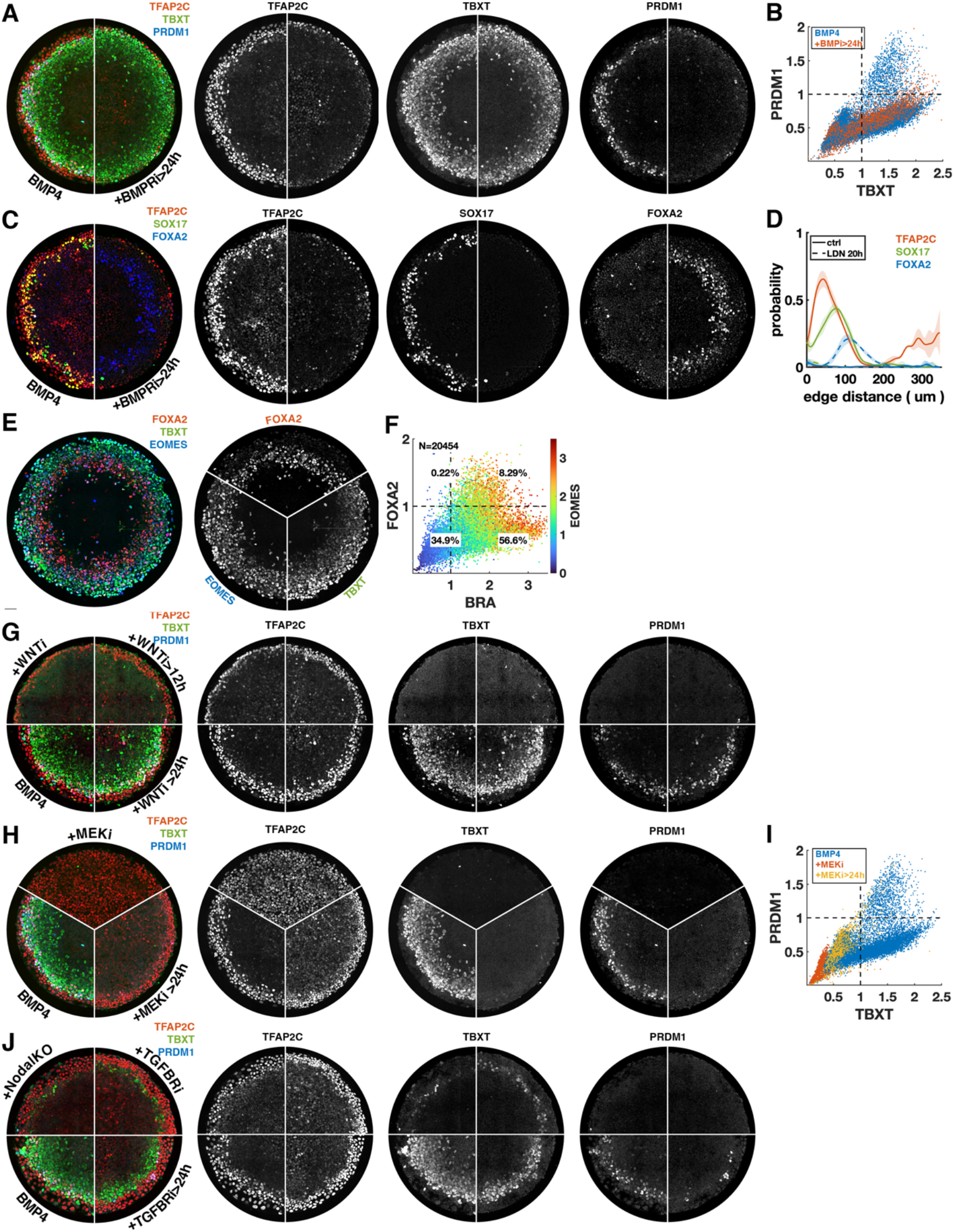
Additional images and quantification for signaling perturbations in figure 3. **a)** Overlay and separate channels for Fig. 3a. **b)** scatterplot of PRDM1 vs. TBXT for BMP4-treated colonies with or without BMP-receptor inhibition after 24h, colored for the condition and **c)** Overlay and separate channels for Fig. 3d and **d)** corresponding radial expression profile. **e-f)** Staining and quantification for FOXA2, TBXT, EOMES of BMP4 treated colony treated with BMPRi after 24h, showing co-expression of FOXA2 and TBXT, which together with lack of SOX17 in c suggests axial mesoderm. **g)** Additional staining for TFAP2C, SOX17,FOXA2 of WNT inhibition after 0 and 24h compared to BMP4 only. **h)** Overlay and separate channels for Fig. 3k. **i)** scatterplot of PRDM1 vs. TBXT for BMP4-treated colonies with or without MEK inhibition after 0 or 24h, colored for the condition. **j)** Overlay and separate channels for Fig. 3p.

**Supplementary figure 6:**
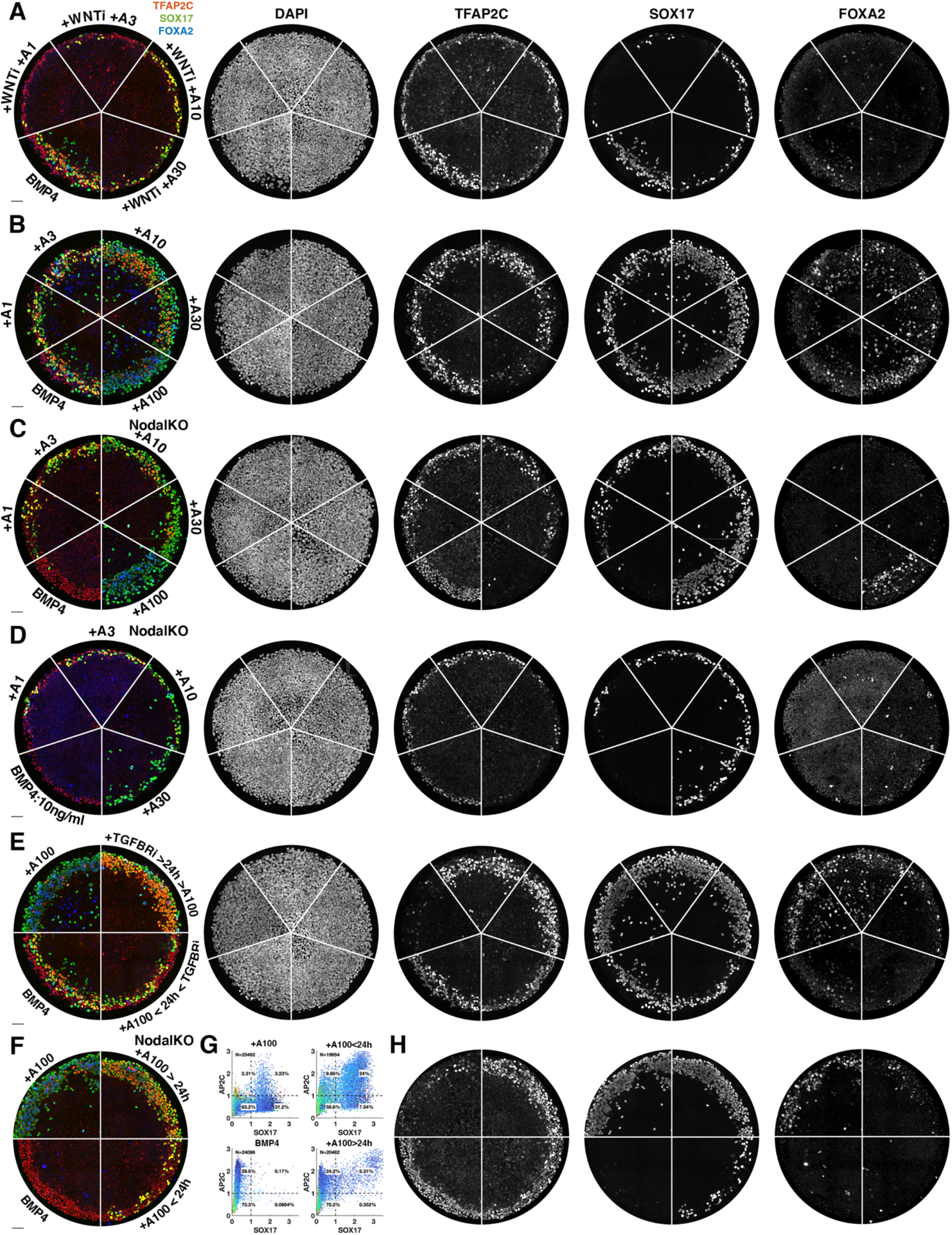
Images of individual channels for figure 4 and effect of Activin timing on NODALKO cells. **a-e)** Images of individual channels for figure 4. a) corresponds to 4a, b) to 4d, c) to 4f, d) to 4j, e) to 4l. **f-h)** Effect of exogenous Activin timing on cell fate in NODALKO cells. **f)** overlay, **g)** quantification, **h)** individual channels.

**Supplementary figure 7:**
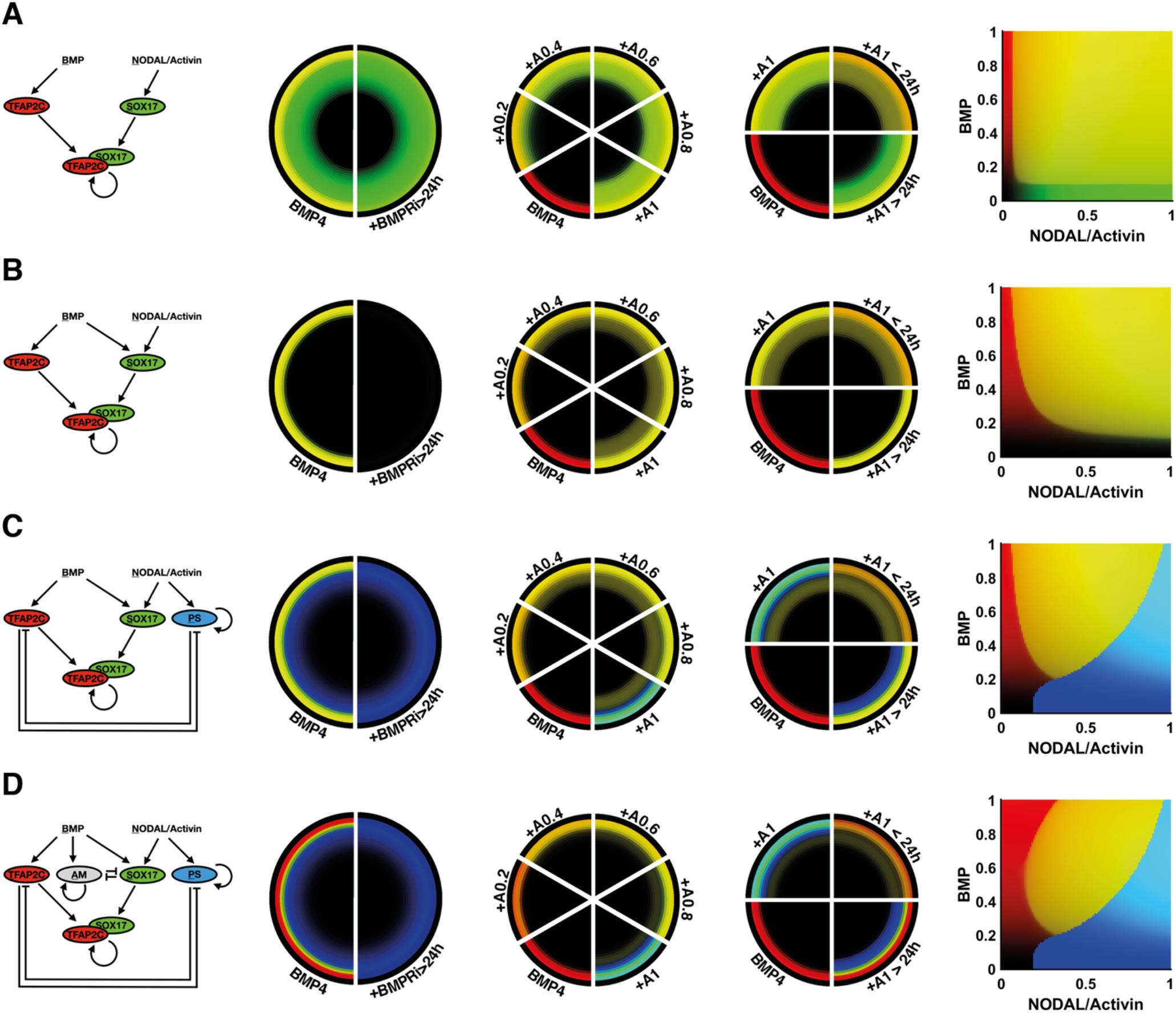
Simpler mathematical models. Each row shows a model and its predicted patterns for the perturbations we wanted to explain. **a)** Stable PGCLCs through autoregulation of SOX17 in combination with TFAP2C and SOX17 activated by NODAL along. **b)** Activation of SOX17 by BMP and NODAL. **c)** Inclusion of a stable PS-like state activated by NODAL which cross-represses with TFAP2C. **d)** Full model including a stable amnion-like state which cross-represses with SOX17.

## SUPPLEMENTARY TABLES

**Supplementary Table 1:** List of relevant genes for gastrulation.

|  |  |  |  |  |  |  |  |
| --- | --- | --- | --- | --- | --- | --- | --- |
| ACVR1 | ACVR1B | ACVR1C | ACVR2A | ACVR2B | ARAF | BMP2 | BMP4 |
| BMP7 | BMPR1A | BMPR2 | BRAF | CDH1 | CDH2 | CDX2 | CER1 |
| COL1A1 | COL6A2 | CXCR4 | DKK1 | DKK3 | DKK4 | DLX5 | DPPA4 |
| DUSP1 | DUSP10 | DUSP14 | DUSP2 | DUSP26 | DUSP3 | DUSP4 | DUSP5 |
| DUSP6 | DUSP7 | DUSP8 | DUSP9 | DVL1 | DVL2 | DVL3 | EGF |
| EGFR | EOMES | EPCAM | EPHA4 | ESRG | ETS1 | EXT1 | EXT2 |
| FGF17 | FGF2 | FGF4 | FGF8 | FGFR1 | FGFR2 | FGFR3 | FGFR4 |
| FGFRL1 | FOXA2 | FZD1 | FZD2 | FZD3 | FZD4 | FZD5 | FZD6 |
| FZD7 | FZD8 | FZD9 | GABRP | GATA2 | GATA3 | GATA4 | GATA6 |
| GLG1 | GPC1 | GPC2 | GPC3 | GPC4 | GPC6 | GSC | HAND1 |
| HEY1 | HHEX | HOXA13 | HOXB2 | HOXB6 | HOXB7 | HOXB9 | HOXD1 |
| HOXD11 | HPSE | HS2ST1 | HS3ST1 | HS3ST3A1 | HS3ST3B1 | HS3ST4 | HS3ST5 |
| HS6ST1 | HS6ST2 | HS6ST3 | HSPG2 | IL17RD | ISL1 | KIT | KLF4 |
| KRT18 | KRT7 | LAMA4 | LEFTY1 | LEFTY2 | LHX1 | LYPD1 | MAP2K1 |
| MAP2K2 | MAPK1 | MAPK3 | MESP1 | MIXL1 | MSGN1 | NANOG | NANOS3 |
| NDST1 | NDST2 | NDST3 | NES | NKX2-5 | NODAL | NOG | OTX2 |
| PAICS | PAIP1 | PAIP2 | PAIP2B | PDGFRA | PEBP1 | PGF | PODXL |
| POSTN | POU5F1 | PRDM1 | RAF1 | RSPO3 | RSPO4 | RUNX1 | SDC1 |
| SDC2 | SDC3 | SDC4 | SMAD1 | SMAD2 | SMAD3 | SMAD4 | SMAD5 |
| SMAD6 | SMAD7 | SMAD9 | SNAI1 | SNAI2 | SOX17 | SOX2 | SOX3 |
| SPRY1 | SPRY2 | SPRY3 | SPRY4 | SULF1 | SULF2 | TBX3 | TBX6 |
| TBXT | TDGF1 | TFAP2A | TFAP2C | TGFBR1 | TGFBR2 | TGFBR3 | TGFBR3L |
| TGFBRAP1 | TNNT2 | TP63 | UGDH | VIM | WNT2B | WNT3 | WNT3A |
| WNT4 | WNT5A | WNT5B | WNT6 | WNT8A | WNT9B | ZIC3 |  |

**Supplementary Table 2:** Top 20 differentially expressed genes between clusters.

|  | iMeLC | pluri | AMLC | PS | PGC | ExMe |
| --- | --- | --- | --- | --- | --- | --- |
| 0 | MIXL1 | SOX2 | ISL1 | GATA6 | SOX17 | SERPINE1 |
| 1 | SERPINE2 | TERF1 | CLDN10 | MESP1 | PRDM1 | NPPB |
| 2 | GAL | ESRG | AMER2 | CER1 | NANOS3 | ANXA1 |
| 3 | MEIS3 | LNCPRESS1 | SGK1 | FGF17 | CRHBP | SPP1 |
| 4 | CNTNAP2 | USP44 | KRT19 | GATA6-AS1 | TFAP2C | MYL7 |
| 5 | EOMES | POLR3G | HPGD | DKK1 | AC007326.5 | TAGLN |
| 6 | CYP1B1 | FOXD3-AS1 | HAPLN1 | MIXL1 | LBH | VIM |
| 7 | WNT3 | DPPA4 | KRT8 | MESP2 | GLIPR2 | ACTA1 |
| 8 | PCDH10 | PLAAT3 | BAMBI | LZTS1 | TMEM64 | FLNC |
| 9 | TBXT | AASS | TMEM54 | CDH2 | PCAT14 | ANKRD1 |
| 10 | IER3 | AL353747.4 | HAND1 | LHX1 | MPC2 | ACTC1 |
| 11 | SP5 | UGP2 | WNT6 | FOXC1 | LINC01198 | CALD1 |
| **12** | GNB4 | RMST | RARRES2 | GSC | LINC00467 | POSTN |
| **13** | PDHB | MIR302CHG | TRIML2 | RSPO3 | CA2 | TAC1 |
| **14** | HEBP2 | PEBP1 | GATA3 | BMP2 | PLPP1 | DIO2 |
| **15** | IFITM1 | TCF7L1 | SEMA3D | CNKSR3 | FAM184A | FOXD1 |
| **16** | NODAL | AC006329.1 | AHNAK | APLNR | NTF3 | IGFBP5 |
| **17** | CDA | SNRPN | EZR | EOMES | AL353807.5 | MIR4435-2HG |
| **18** | CCND2 | SEPHS1 | RHOBTB3 | ARL4D | PRTG | CCN2 |
| **19** | KCNG1 | VSNL1 | EFNA1 | SNAI1 | DBI | CCN1 |

**Supplementary Table 3:** Top 20 differentially expressed genes between clusters within set of genes relevant for gastrulation.

|  | iMeLC | pluri | AMLC | PS | PGC | ExMe |
| --- | --- | --- | --- | --- | --- | --- |
| **0** | MIXL1 | SOX2 | ISL1 | GATA6 | SOX17 | VIM |
| **1** | EOMES | ESRG | HAND1 | MESP1 | PRDM1 | POSTN |
| **2** | WNT3 | DPPA4 | WNT6 | CER1 | NANOS3 | COL1A1 |
| **3** | TBXT | PEBP1 | GATA3 | FGF17 | TFAP2C | SOX3 |
| **4** | NODAL | TDGF1 | BMP4 | DKK1 | TFAP2A | NES |
| **5** | FGF4 | LEFTY1 | DLX5 | MIXL1 | WNT5B | DUSP8 |
| **6** | FGF8 | NANOG | KRT18 | CDH2 | TBX3 | PODXL |
| **7** | CDH2 | PODXL | TFAP2A | LHX1 | CXCR4 | NOG |
| **8** | IL17RD | SPRY1 | TBX3 | GSC | GATA3 | FZD2 |
| **9** | DUSP5 | FZD5 | EPCAM | RSPO3 | VIM | NKX2-5 |
| **10** | TDGF1 | LEFTY2 | KRT7 | BMP2 | DKK3 | SULF2 |
| **11** | WNT8A | FGFR2 | WNT5B | EOMES | CDX2 | KRT18 |
| **12** | GSC | FGFR1 | HOXB2 | SNAI1 | ACVR1 | GPC4 |
| **13** | DUSP26 | GPC4 | DUSP2 | PDGFRA | HS3ST5 | TFAP2C |
| **14** | PEBP1 | POU5F1 | NOG | TBXT | SMAD9 | HSPG2 |
| **15** | COL6A2 | PAICS | HOXB6 | EPHA4 | KRT18 | CDH2 |
| **16** | GPC6 | FGF2 | HEY1 | OTX2 | ISL1 | GATA3 |
| **17** | FGF17 | DUSP14 | TFAP2C | WNT5A | SPRY4 | SDC3 |
| **18** | NDST3 | NODAL | LYPD1 | FGF4 | MESP1 | SDC2 |
| **19** | PAICS | ZIC3 | SMAD9 | DKK4 | SDC2 | DKK3 |

## Notes

### Competing Interest Statement

The authors have declared no competing interest.

