## Supplementary Text on Image Analysis for "Efficient differentiation of human primordial germ cells through geometric control reveals a key role for NODAL signaling"

### 3D segmentation algorithm

Nuclei were segmented in individual z-slices based on DAPI staining using two different freely available machine learning-based software tools: Ilastik (Berg et al. 2019) and Cellpose (Stringer et al. 2021). We found that Cellpose accurately determines masks for individual nuclei and outputs them as a label matrix, but has a high false negative rate. Ilastik, on the other hand, generates a binary mask of foreground versus background that includes all nuclei in the image but does not separate them. Because Cellpose generally correctly identifies all of the masks for the nuclei that it detects, we first segment each z slice of the nuclear channel with Cellpose, and we independently segment the image into foreground and background with Ilastik. To fill in the gaps in the Cellpose segmentation, we take all pixels of the Ilastik foreground mask that are not members of any Cellpose nuclear mask as a binary mask of all the nuclei missed by Cellpose. We remove small noise features from this mask with a morphological opening operation and identify connected components of the resulting mask as individual nuclei at each z slice, separating merged or overlapping nuclei with a convex decomposition algorithm based on (Lien and Amato 2004).

To determine the links between nuclei in adjacent frames, we used the framework of a linear assignment problem (LAP), based loosely on the approach to single-particle tracking in Jaqaman et al. 2008. To link nuclei in slice $z_n$ to nuclei in slice $z_{n+1}$, we defined the cost matrix for the LAP as a block matrix of the form

$$\begin{bmatrix} A & B \\ C & A^T \end{bmatrix}.$$

$A(i,j)$ gives the cost of linking nucleus $i$ in frame $n$ to nucleus $j$ in frames $n+1$, and is given by

$$A(i,j) = \begin{cases} \frac{\min(|N_{n,i}|,|N_{n+1,j}|)}{|N_{n,i} \cap N_{n+1,j}|} & \text{if } d(i,j) \leq d_{\max} \\ \text{Inf} & \text{if } d(i,j) > d_{\max} \end{cases},$$

where $N_{n,i}$ is the set of pixels in the mask of nucleus $i$ in frame $j$ and $d(i,j)$ is the distance between the centroids of the two masks. That is, the cost to link two nuclei is the smaller of the sizes of the two masks divided by the size of their overlap. For efficiency, each nucleus in slice $n$ has this cost computed only for its three nearest neighbors in slice $n+1$ and vice versa, and all other costs are set to Inf (arbitrarily large, so that these links are treated as impossible). We further impose a cutoff $d_{\max}$ on the distance between the centroids of the two nuclei and set $A(i,j) = \text{Inf}$ if the distance exceeds the cutoff. Finally, $B$ and $C$ are square diagonal matrices with all off-diagonal entries set to Inf and diagonal entries set to the "alternative cost" 1/IoU for not linking to any other nucleus, where IoU is an intersection over union threshold set to determine the minimum ratio of overlap to nucleus area that qualifies two nuclei to be linked. If every cost along the $i^{\text{th}}$ row of $A$ exceeds 1/IoU, then nucleus $i$ in slice $n$ will be linked to nothing, and likewise for costs along columns.

This linking operation is performed sequentially across pairs of adjacent z slices, creating chains of linked masks in different slices that are taken to correspond to a single nucleus. We additionally impose a maximum expected nuclear diameter and use the spacing between z slices to determine the maximum number of slices that may correspond to a single nucleus. If more than this number of masks are linked together, the chain is broken into two parts by splitting it at a local minimum in the area of the nuclear mask. Since nuclei are defined across multiple z slices, a given nucleus has a readout of average fluorescent intensity in each channel in each slice. For each channel we take the maximum across z slices as the value for that nucleus, as it should correspond to the readout in which the nucleus was most nearly in focus.
