## Supplementary Text on Mathematical Model for "Efficient differentiation of human primordial germ cells through geometric control reveals a key role for NODAL signaling"

### ODE model for hPGCLC specification

**Desired qualitative behavior**

We aimed to identify the minimal mathematical model that recapitulates the following qualitative features of PGCLC specification in response to BMP and NODAL/Activin signaling:

1. TFAP2C does not depend on NODAL but is eliminated by adding BMPRi after 24h, while SOX17 does require NODAL (Fig 3A,P).

2. PGCs are stable and express higher TFAP2C than TFAP2C+SOX17- cells which disappear over time (Fig 1,2).

3. TFAP2C and SOX17 are expressed in concentric rings that never extend into the colony much more than 100um (Fig 1A).

4. High Activin throughout differentiation eliminates TFAP2C expression and gives rise to mesoderm and endoderm while lower doses have only a small effect in WT cells and rescue WT levels in NodalKO with outer cells remaining SOX17- (Fig 4D-G,L-O).

5. High Activin during only the first 24h results in SOX17 coexpression in all TFAP2C+ cells, while high Activin only after the first 24h leads to much less SOX17 with outer cells remaining TFAP2C+/SOX17- (Fig 4L-O).

**GRN construction**

To explain this set of observations, we formulated a hypothetical gene regulatory network (GRN) integrating NODAL/Activin and BMP signaling to decide between amnion-like, mesendodermal, and primordial germ cell-like fates. For simplicity, all activating and inhibitory interactions are taken to act transcriptionally rather than as post-transcriptional or post-translational interactions.

To explain the first observation, and in agreement with Kojima, Sasaki, et al. 2017 we take TFAP2C to be directly activated by BMP signaling and take SOX17 to be activated by Activin/NODAL signaling. SOX17 induction also depends on NODAL indirectly through EOMES but we not include this complication in our model for simplicity.

The second observation suggests that expression of TFAP2C and SOX17 is maintained by positive feedback, but that this positive autoregulation functions only when both genes are present. Several mechanisms could explain this behavior; for instance, SOX17 and TFAP2C could bind independently at different regions of their promoters, but stimulate transcription only when both are present. Alternatively, SOX17 and TFAP2C may only regulate their own transcription in complex with one another. Because SOX proteins are known to complex with a partner transcription factor (TF) for their action (Kamachi and Kondoh 2013), we favor the latter explanation and model SOX17 and TFAP2C as forming a dimer to upregulate their own expression but this is not essential for the behavior of the model. This results in a stable population of SOX17+/TFAP2C+ cells, but no stable SOX17-/TFAP2C+ population.

The third observation requires a mechanism to restrict expression of both genes to a ring near the colony edge in all cases. It has been shown (Etoc et al. 2016; Heemskerk et al. 2019; Chhabra et al. 2019) that in BMP4-driven differentiation of micropatterned hPSC colonies, BMP signaling is initially uniformly high throughout the colony, before becoming restricted to the colony edge with a sharp gradient towards the colony center after about 12 hours due to receptor relocalization and production of diffusible BMP inhibitors (Fig. 6A). We expect that TFAP2C expression is restricted to near the colony edge because this is the only region of the colony that experiences sustained high BMP signaling. However, in wild type (WT) micropatterned colonies, it has been observed (Heemskerk et al. 2019; Chhabra et al. 2019) that a wave of high NODAL signaling begins at the colony edge at about 24 hours and moves inward towards the colony center at a constant velocity, reaching most of the colony by 42 hours, so that the range of SOX17 expression cannot be explained only by the range of NODAL signaling (Fig. 6A). We propose that SOX17 requires both NODAL and BMP signaling at relatively low levels in order to restrict expression from the colony center. An alternative explanation could be that instead of relying directly on BMP, SOX17 relies on induction by a BMP target TF such as TFAP2C as suggested by Chen et al. 2019, or GATA3 as suggested by Kojima, Yamashiro, et al. 2021.

The center of wild type BMP4-treated micropatterns experiences high BMP signaling only during the first 12 hours of differentiation and high NODAL signaling only after the first 24, preventing induction of SOX17 in this region in our model. Because SOX17 is absent in the colony center, any initial expression of TFAP2C in response to high BMP signaling in this region is not sustained. Also note that it has been observed (Etoc et al. 2016) that densely-packed hPSCs have reduced sensitivity to (apical) Activin treatment due to receptor relocalization, so that Activin-treated colonies form a gradient of Smad2/3 activity that is highest at the colony edge, with low signaling in the center. However, we expect the gradient from the colony edge to be less steep than for BMP since, unlike for BMP, there is no evidence of a spatial profile of signaling inhibitors that is highest in the colony center (Fig. 6A).

Observation four suggests that Activin can inhibit TFAP2C expression, but only at the highest dose and for treatment that is sustained for >24 hours. A simple mechanism that could explain this observation is that Activin stimulates the production of another TF that inhibits TFAP2C but only after it accumulates to a sufficiently high level. Furthermore, for the highest Activin rescue dose in both WT and NODALKO cells, there remains a ring of SOX17 expression of similar size, but most of these cells coexpress the definitive endoderm gene FOXA2 instead of the PGC marker TFAP2C, indicating that differentiation is switched to mesendoderm. For this reason, we take the proposed inhibitory TF acting on TFAP2C to be a specifier of primitive streak and refer to it generically as PS. We further observed that inhibition of TFAP2C, once it does happen, appears to be complete, so that high sustained Activin fully represses TFAP2C, while lower or transient treatment has little or no effect. To explain both the observed delay before inhibition begins, and its switch-like nature, we propose that PS initially slowly accumulates in response to NODAL/Activin signaling before reaching a threshold for autoactivation, after which it rapidly stimulates its own production to a high level. A similar separation of production into an initial slow regime followed by fast autoactivation was suggested by the transcriptional dynamics of a subset of BMP4 and Activin target genes in Heemskerk et al. 2019, including the primitive streak markers TBXT and EOMES. An alternative mechanism to explain this delay and switch-like activation could be an additional TF upstream of PS that activates it in a coherent feedforward loop with Activin signaling; to keep the minimal number of genes in the network, we model only the first proposed mechanism. We additionally take TFAP2C to repress production of PS in agreement with the observation in Chen et al. 2019 that TFAP2C inhibits differentiation to primitive streak derivatives, and to explain the observation that higher BMP doses require a higher Activin dose to repress PGCLC fate (Fig 4).

The final observation suggests that sustained high BMP signaling at the outermost edge of the colony in the absence of NODAL/Activin signaling causes cells to commit to an amnion-like fate that prevents later induction of SOX17 in response to Activin or NODAL. To impose this behavior, we included an amnion-specific TF, referred to as AM, with a formulation similar to that of PS: slow accumulation in response to high BMP signaling before reaching an autoactivation threshold and robustly upregulating its own production, and we also take AM and SOX17 to be mutually inhibitory. Because we see that the outermost cells in the colony become TFAP2C+/SOX17- and a ring inset from these becomes TFAP2C+/SOX17+, AM autoactivation must occur early enough to prevent induction of SOX17 only with the highest levels of sustained BMP at the colony edge, while TFAP2C and SOX17 can respond to intermediate levels farther down the BMP gradient. An alternative and possibly redundant mechanism to prevent PGCLC differentiation on the colony edge could be BMP-depedent desensitization to NODAL/Activin signaling via downregulation of TGF$\beta$ receptors and upregulation of NODAL/Activin inhibitors; for simplicity we did not include this in the model.

We tested systematically if the model identified this way could be further simplified by taking parts out and testing the predicted cell fate patterns for each perturbation (SI Fig. 7). Each of the simplified models failed to correctly predict part of the patterns.

**ODE model**

Denote NODAL/Activin as $N$, BMP as $B$, TFAP2C as $T$, SOX17 as $S$, and the primitive streak and amnion transcription factors PS and AM as $P$ and $A$, respectively. Degradation rates are denoted $\alpha$, maximal production rates are denoted $\beta$, and $K_{XY}$ gives the activation or inhibition threshold for $X$ acting on $Y$. Each transciption factor is assumed to positively regulate its own production, and all input functions are taken as Hill functions with coefficient $n$. We also take the combined dilution and degradation rate $\alpha$ for each gene to be approximately equal. In agreement with the simple GRN sketched above, we can write the following system of ordinary differentiation equations (ODEs):

$$\frac{dT}{dt} = \beta_T \frac{(B/K_{BT})^n + \big([\mathrm{TS}]/K_{[\mathrm{TS}]}\big)^n}{1 + (B/K_{BT})^n + \big([\mathrm{TS}]/K_{[\mathrm{TS}]}\big)^n} \cdot \frac{K_{PT}^n}{K_{PT}^n + P^n} - \alpha T, \tag{1}$$

$$\frac{dS}{dt} = \beta_S \frac{(NB/K_{NBS})^n + \big([\mathrm{TS}]/K_{[\mathrm{TS}]}\big)^n}{1 + (NB/K_{NBS})^n + \big([\mathrm{TS}]/K_{[\mathrm{TS}]}\big)^n} \cdot \frac{K_{AS}^n}{K_{AS}^n + A^n} - \alpha S, \tag{2}$$

$$\frac{dP}{dt} = \left(\frac{\beta_{NP} N^n}{K_{NP}^n + N^n} + \frac{\beta_{PP} P^n}{K_{PP}^n + P^n}\right) \cdot \frac{K_{TP}^n}{K_{TP}^n + T^n} - \alpha P, \tag{3}$$

$$\frac{dA}{dt} = \left(\frac{\beta_{BA} B^n}{K_{BA}^n + B^n} + \frac{\beta_{AA} A^n}{K_{AA}^n + A^n}\right) \cdot \frac{K_{SA}^n}{K_{SA}^n + S^n} - \alpha A, \tag{4}$$

$$[\mathrm{TS}] = K_d T_{\mathrm{free}} S_{\mathrm{free}}, \qquad T_{\mathrm{free}} = T - [\mathrm{TS}], \qquad S_{\mathrm{free}} = S - [\mathrm{TS}]. \tag{5}$$

The concentration of the dimer [TS] in (5) is determined by the equation

$$\frac{d[\mathrm{TS}]}{dt} = \gamma_{TS} T_{\mathrm{free}} S_{\mathrm{free}} - \gamma_{[\mathrm{TS}]}[\mathrm{TS}], \tag{6}$$

where $\gamma_{TS}$ and $\gamma_{[\mathrm{TS}]}$ are association and disassociation constants for the complex. Since complex formation happens much faster than protein production and degradation, we can take this equation to be effectively at equilibrium on the timescale of the above system of ODEs, so that

$$[\mathrm{TS}] = K_d T_{\mathrm{free}} S_{\mathrm{free}}, \tag{7}$$

where $K_d = \gamma_{TS}/\gamma_{[\mathrm{TS}]}$. We can express $T_{\mathrm{free}}$ and $S_{\mathrm{free}}$ in terms of the total concentration of $T$ and $S$ as $T_{\mathrm{free}} = T - [\mathrm{TS}]$ and $S_{\mathrm{free}} = S - [\mathrm{TS}]$. Substituting into (7) and rearranging yields

$$[\mathrm{TS}]^2 - (T + S + 1/K_d)[\mathrm{TS}] + TS = 0,$$

which has the solutions

$$[\mathrm{TS}] = \frac{T + S + 1/K_d \pm \sqrt{(T + S + 1/K_d)^2 - 4TS}}{2}.$$

Given the additional constraint that $[\mathrm{TS}] \leq \min(T, S)$, we can discard the larger root as it is always larger than both $T$ and $S$, so that

$$[\mathrm{TS}] = \frac{T + S + 1/K_d - \sqrt{(T + S + 1/K_d)^2 - 4TS}}{2}.$$

Note that as $K_d$ becomes large, [TS] approaches $\min(T, S)$. When numerically evaluating the system of ODEs, the dimer concentration is set to this steady-state value depending on absolute concentrations of $T$ and $S$ at each step.

In (1) and (2), $T$ and $S$ have activation functions written such that production by either autoactivation or signaling inputs is a Hill function with coefficient $n$, but with the Hill functions combined to approximate OR logic; that is, for either $[\mathrm{TS}] \gg K_{[\mathrm{TS}]}$ or $B \gg K_{BT}$,

$$\beta_T \frac{(B/K_{BT})^n + \left([\mathrm{TS}]/K_{[\mathrm{TS}]}\right)^n}{1 + (B/K_{BT})^n + \left([\mathrm{TS}]/K_{[\mathrm{TS}]}\right)^n} \to \beta_T,$$

so that production of $T$ can be driven at similar levels by either $B$ or [TS]. Similarly, production of $S$ can be driven either by [TS] or by $A$ and $B$ together. The inhibition function for each gene, on the other hand, is combined multiplicatively to approximate AND logic so that sufficiently high levels of inhibitor can completely downregulate production. For instance, $T$ is produced only if ($[\mathrm{TS}] > K_{[\mathrm{TS}]}$ OR $B > K_{BT}$) AND ($P < K_{PT}$).

Equations (1) and (2) allow a relatively short exposure time to BMP and Activin to induce TFAP2C and SOX17. We wrote (3), however, according to the observation that PS turns on only after a substantial delay, and is then rapidly upregulated to a high level to completely repress TFAP2C. We implemented this behavior by making $P$ be activated at a low level by $A$, and at a much higher level by autoactivation by making the production terms for $P$ additive with $\beta_{PP} > \beta_{NP}$. This separates production of $P$ into an initial slow regime of Activin-driven production before reaching a threshold level for autoactivation and then accumulating much faster, as described in the **GRN construction** section. In the initial regime, assuming no inhibition from $T$, we have

$$\frac{dP}{dt} = \frac{\beta_{NP} N^n}{K_{NP}^n + N^n} - \alpha P, \tag{8}$$

so that if $N > K_{NP}$, $P$ accumulates towards a steady-state level of $\mathrm{ss}_{low} = \beta_{NP}/\alpha$ before reaching its autoactivation threshold. Once the threshold has been reached, if Activin is removed, $P$ accumulates according to

$$\frac{dP}{dt} = \frac{\beta_{PP} P^n}{K_{PP}^n + P^n} - \alpha P. \tag{9}$$

We find the autoactivation-driven steady state by setting $dP/dt = 0$ in (9). Doing this and rearranging, we get

$$P^{n+1} - \frac{\beta_{PP}}{\alpha} P^n + K_{PP}^n P = 0.$$

Letting $n = 2$, we have

$$P\left(P^2 - \frac{\beta_{PP}}{\alpha}P - K_{PP}^2\right) = 0,$$

which has the solutions $P = 0$ and

$$P = \frac{\beta_{PP}/\alpha \pm \sqrt{(\beta_{PP}/\alpha)^2 - 4K_{PP}^2}}{2},$$

so that

$$P = 0, \quad P = \frac{1}{2}\beta_{PP}/\alpha + \frac{1}{2}\sqrt{(\beta_{PP}/\alpha)^2 - 4K_{PP}^2}$$

are stable fixed points and

$$P = \frac{1}{2}\beta_{PP}/\alpha - \frac{1}{2}\sqrt{(\beta_{PP}/\alpha)^2 - 4K_{PP}^2}$$

is an unstable fixed point between them. Then, following the above analysis, the smaller non-zero root is the threshold, $\tau_P$, for autoactivation of P, and the larger root is the autoactivation-driven steady state, $\text{ss}_{high}$. Setting $\tau_P$ and $\text{ss}_{high}$ to desired values, we can write $K_{PP} = \sqrt{\text{ss}_{high}\tau_P}$ and $\beta_{PP} = \alpha(\text{ss}_{high} + \tau_P)$ in (3). In the case that the autoactivation threshold has been met and signaling remains on, the steady state level reached will be $\text{ss}_{high} + \text{ss}_{low}$, but for $\text{ss}_{high} \gg \text{ss}_{low}$, we can take this to be approximately $\text{ss}_{high}$. Because we also expected $A$ to initially accumulate slowly before being robustly autoactivated, we wrote the activation function of $A$ with the same structure as $P$, so that the analysis for autoactivation of $A$ is identical to that for $P$.

**Parameter considerations**

For simplicity, and in the absence of concrete data on protein production and degradation rates, we set $\alpha_T = \alpha_S = \alpha_P = \alpha_A = \alpha$, taking each protein to degrade and dilute at approximately the same rate (and in fact, if we take the proteins to be stable the term $\alpha$ in each equation only describes dilution due to cell growth and division, which is constant across all proteins). Because the units of protein concentration in the simulation are arbitrary, we set $\beta_T = \beta_S = \alpha$ and $\text{ss}_{high} = 1$ for both $P$ and $A$, so all concentrations vary between 0 and 1. We can additionally express the $\beta$ and $K$ parameters for $P$ and $A$ in terms of thresholds and steady states as

$$\begin{aligned}
\beta_{NP} &= \alpha \cdot \text{ss}_{lowP} \\
\beta_{BA} &= \alpha \cdot \text{ss}_{lowA} \\
K_{PP} &= \sqrt{\text{ss}_{high} \cdot \tau_P} = \sqrt{\tau_P} \\
K_{AA} &= \sqrt{\text{ss}_{high} \cdot \tau_A} = \sqrt{\tau_A} \\
\beta_{PP} &= \alpha(\text{ss}_{high} + \tau_P) = \alpha(1 + \tau_P) \\
\beta_{AA} &= \alpha(\text{ss}_{high} + \tau_A) = \alpha(1 + \tau_A)
\end{aligned}$$

Because system behavior is similar for a reasonable range of $n$ from 2 to 4, we let it be the same for each equation and set $n = 2$. Then we can rewrite (1) through (4) as

$$\frac{1}{\alpha}\frac{dT}{dt} = \frac{(B/K_{BT})^2 + \left([\mathrm{TS}]/K_{[\mathrm{TS}]}\right)^2}{1 + (B/K_{BT})^2 + \left([\mathrm{TS}]/K_{[\mathrm{TS}]}\right)^2} \cdot \frac{K_{PT}^2}{K_{PT}^2 + P^2} - T \tag{10}$$

$$\frac{1}{\alpha}\frac{dS}{dt} = \frac{(NB/K_{NBS})^2 + \left([\mathrm{TS}]/K_{[\mathrm{TS}]}\right)^2}{1 + (NB/K_{NBS})^2 + \left([\mathrm{TS}]/K_{[\mathrm{TS}]}\right)^2} \cdot \frac{K_{AS}^n}{K_{AS}^2 + A^n} - S \tag{11}$$

$$\frac{1}{\alpha}\frac{dP}{dt} = \left(\frac{\mathrm{ss}_{lowP} N^2}{K_{NP}^2 + N^2} + \frac{(1+\tau_P)P^2}{\tau_P + P^2}\right) \cdot \frac{K_{TP}^2}{K_{TP}^2 + T^2} - P \tag{12}$$

$$\frac{1}{\alpha}\frac{dA}{dt} = \left(\frac{\mathrm{ss}_{lowA} B^2}{K_{BA}^2 + B^2} + \frac{(1+\tau_A)A^2}{\tau_A + A^2}\right) \cdot \frac{K_{SA}^2}{K_{SA}^2 + S^2} - A \tag{13}$$

We also take BMP and Activin/NODAL activity to vary between minimum and maximum values of 0 and 1, and set activation thresholds in this range. Take $K_d$ to be large (initially $K_d = 100$), so that most available $X$ and $Y$ dimerize to form [XY]; note that the autoactivation threshold $K_{[\mathrm{XY}]}$ can be tuned to compensate for different values of $K_d$. The expected qualitative behavior of the model now mainly depends on thresholds for autoactivation and the thresholds for inhibition between $T$ and $P$ and between $S$ and $A$, which can be tuned to reasonable ranges.

Finally, note that in our simulations we take BMP and NODAL-driven gene activation to rely on (sigmoidal) Hill functions where AM has a higher BMP threshold for activation than TFAP2C and PS has a higher NODAL threshold for activation than SOX17. This was a natural way to explain the observation that TFAP2C+ cells generally expand farther into the colony than the AmLC fate ring, and that SOX17 is expressed with very low Activin rescue doses while TFAP2C is only repressed by the proposed PS gene for the highest Activin dose. However, this is not an essential feature of the model, and similar qualitative behavior can be obtained if BMP and NODAL activate gene expression with first-order (non-sigmoidal) Hill functions by tuning the relative inhibition strengths between TFAP2C and PS and between SOX17 and AM, as well as each gene's threshold for autoactivation.

Referring to the analysis of how a delay is imposed on robust activation of PS, recall that, neglecting inhibition, NODAL/Activin-mediated accumulation of $P$ follows (8), so that for a given signaling input level $N$, it approaches the steady state

$$ss_{\mathrm{low}} = \frac{\beta_{NP}}{\alpha} \cdot \frac{N^n}{K_{NP}^n + N^n}.$$

If activation by signaling is switch-like (high $n$), this can take on values of 0 or $\beta_{NP}/\alpha$, and whether the autoactivation threshold can be reached depends on whether the signaling input is above the threshold $K_{NP}$. If signaling-driven production is taken to be more graded (low $n$), then the steady-state can take on a range of values between 0 and $\beta_{NP}/\alpha$. In this case, there is still a specific level of NODAL/Activin above which it is possible to reach the autoactivation threshold, but how long it takes to reach the threshold depends the level of $N$. The signaling-mediated steady state is further lowered by the presence of inhibitors, which explains, for instance, why high BMP for the first 24 hours in the absence of NODAL/Activin is taken to sufficiently induce production of AM to a high level, but if BMP and Activin are concurrently applied, SOX17 outcompetes AM, except at the colony edge at the lowest rescue dose. Further experimental work and analysis of the robustness of the system to perturbations in parameter values will shed light on which of these scenarios is more plausible.

To generate the results shown in Fig. 6 and SI Fig. 7I, we used the following parameter values:

| Parameter | Value | Meaning |
|---|---|---|
| n | 2 | Hill function coefficient |
| $\alpha$ | 0.1733 | protein dilution + degradation rate |
| $(\beta_T, \beta_S)$ | 0.1733 | production rate for $T$ and $S$ |
| $\text{ss}_{high}$ | 1 | autoactivation steady state for $P$ and $A$ |
| $(K_{BT}, K_{NBS}, K_{NZ}, K_{BA})$ | (0.3, 0.18, 0.8, 0.95) | signaling thresholds |
| $(K_{PT}, K_{AS}, K_{TP}, K_{SA})$ | $(0.1, 0.08, 0.0.4, 0.85)$ | inhibition thresholds |
| $(\text{ss}_{lowP}, \text{ss}_{lowA})$ | (0.2, 0.1) | maximum signaling-driven protein level |
| $(\tau_P, \tau_A)$ | $(0.04, 0.12)$ | autoactivation thresholds |
| $K_{[\text{TS}]}$ | 0.5 | activation threshold for [TS] on $T$ and $S$ |
| $K_d$ | 100 | dimerization constant for [TS] |
